# Multiplexed action-outcome representation by striatal striosome-matrix compartments detected with a novel cost-benefit foraging task

**DOI:** 10.1101/2021.08.17.456542

**Authors:** Bernard Bloem, Rafiq Huda, Ken-ichi Amemori, Alexander Abate, Gaya Krishna, Anna Wilson, Cody W. Carter, Mriganka Sur, Ann M. Graybiel

**Author notes:** These authors contributed equally to this work (Bernard Bloem, Rafiq Huda).

## Abstract

Learning about positive and negative outcomes of actions is crucial for survival and underpinned by conserved circuits including the striatum. How associations between actions and outcomes are formed is not fully understood, particularly when the outcomes have mixed positive and negative features. We developed a novel foraging (‘bandit’) task requiring mice to maximize rewards while minimizing punishments. By 2-photon Ca^++^ imaging, we monitored activity of 5831 identified anterodorsal striatal striosomal and matrix neurons. Surprisingly, we found that action-outcome associations for reward and punishment were combinatorially encoded rather than being integrated as overall outcome value. Single neurons could, for one action, encode outcomes of opposing valence. Striosome compartments consistently exhibited stronger representations of reinforcement outcomes than matrix, especially for high reward or punishment prediction errors. These findings demonstrate a remarkable multiplexing of action-outcome contingencies by single identified striatal neurons and suggest that striosomal neurons are differentially important in action-outcome learning.

## INTRODUCTION

Behavior is powerfully sculpted by learning from reinforcement, with rewards increasing and punishments decreasing the propensity to engage in specific actions (Sutton & Barto, 1998). The striatum has repeatedly been implicated in reinforcement learning (RL) mechanisms that allow animals to adapt their behavior in changing environments by monitoring associations between actions and outcomes (Adams & Dickinson, 1981; Gremel & Costa, 2013; O’Doherty et al., 2004; Simon et al., 2015; Smith & Graybiel, 2016; Thorn et al., 2010; Yang & Masmanidis, 2020; Yin et al., 2005, 2009). Striatal projection neurons (SPNs) encode associations between actions and rewarding outcome, i.e., outcome activity is specific for actions (Lau & Glimcher, 2007; Stalnaker et al., 2010; Thorn et al., 2010). However, in naturalistic settings, the same action could produce both rewarding and aversive outcomes (Amemori & Graybiel, 2012; Aupperle & Paulus, 2010; Friedman et al., 2015; Wallis & Rushworth, 2014). Extensive research applying activity manipulation techniques has shown that aversive learning also depends on the striatum (Delgado et al., 2008; Hikida et al., 2010; Kravitz et al., 2012; Palminteri et al., 2012; Stephenson-Jones et al., 2020). Therefore, a fundamental remaining question is how SPNs represent opposing reinforcement consequences of actions and use them for learning.

The underlying mechanisms of RL are often studied using probabilistic and non-stationary bandit tasks, which require trial-and-error learning in order to maximize only rewards to be obtained (Averbeck & Costa, 2017; Hamid et al., 2015; Hattori et al., 2019; Lau & Glimcher, 2008; Neftci & Averbeck, 2019; Nonomura et al., 2018; Parker et al., 2016; Samejima et al., 2005; Sugrue et al., 2004; Sutton & Barto, 1998; Tai et al., 2012). We developed a ‘cost-benefit bandit (CBB) task’ to bring the advantages of bandit task protocols to address this mixed outcome context. In this dynamic foraging task, each of two available actions is probabilistically linked to outcomes of opposing valence, and the mice learn action-outcome contingencies to maximize reward delivery and minimize air puff delivery. We use here the term action-outcome contingency to highlight the relationship between actions and the outcomes that are associated with these actions.

Although the striatum has been implicated in both appetitive and aversive learning, it is currently unclear whether this system learns rewarding and aversive outcomes of actions in parallel, or whether it learns the overall value of actions. We first asked whether the activity of SPNs represents integrated values of actions or separately represents each element of the outcome. We recorded neuronal activity of thousands of SPNs during the CBB task by 2-photon Ca^++^ imaging and tracked their encoding of action-reward-airpuff associations in the mid-dorsal sector of the caudatoputamen. We could then examine whether, and how, they could encode associations between actions and both rewarding and aversive outcomes. We found large numbers of SPNs responding in relation to both rewarding and aversive outcomes of a given action. To our surprise, among all outcome-responsive SPNs, equal numbers of neurons were active in relation to outcomes with the same or opposite valence. On a population level, both outcomes were represented independently in two partially overlapping populations of neuron in a multiplexed manners. This finding is important, as it suggests that striatal reward-related activity might not reflect the integrated value, but rather, specific outcomes.

The RL perspective hypothesizes that the value of actions is updated using prediction errors (PEs). The striatum has been implicated in the PE signals (Dahlin et al., 2008; Matamales et al., 2020), which are transferred to dopamine-related circuitry (Cohen et al., 2012; Menegas et al., 2018; Schultz et al., 1997; Schultz, 2016; Steinberg et al., 2013). However, it is still unclear whether SPNs represent the PEs by integrating the outcomes of opposing valence, or independently represent reward prediction errors (RPEs) and punishment prediction errors (PPEs). We examined the SPN’s representation of PEs that were inferred from two different RL models. In the first model, agents learn action values by the PEs derived by integrating rewarding and aversive values. In the other model, rewarding and aversive outcomes of available actions were estimated in parallel, and these expectations were integrated when decisions are made. Therefore, the first model calculated one integrated PE, and the other model calculated RPE and PPE independently. The activity of outcome-selective neurons could much better be explained by separate RPE and PPE as compared to a single integrated PEs. This finding further emphasizes the parallel coding of rewarding and aversive learning variables in the striatum.

Computational work further suggested that SPNs in the striosome compartment might specifically function in shaping the PE signals, thereby providing learning signals for matrix neurons to learn action values (Brown et al., 1999; Doya, 2000; Houk et al., 1995; Joel et al., 2002; Takahashi et al., 2009). It is thus possible that the PE-based updating, and modulation of dopaminergic activity, could arise, in part at least, from striosomes, based on anatomical and electrophysiological studies demonstrating strong projections from the striosome compartment to dopamine-containing neurons in the substantia nigra, pars compacta (Crittenden et al., 2016; Evans et al., 2019; Fujiyama et al., 2011; Matsushima & Graybiel, 2020; McGregor et al., 2019).

The use of approach-avoidance paradigms has suggested that striosomal circuits could be part of a conserved mechanism for facing critical decisions requiring an estimate of cost-benefit evaluation (Amemori & Graybiel, 2012; Amemori et al., 2020; Friedman et al., 2015, 2017, 2020). To address this issue, we visually identified SPNs in the most dorsal band of striosomes (sSPNs) and SPNs in the surrounding matrix (mSPNs) by using their birthdates to label striosomes (Bloem et al., 2017; Kelly et al., 2018; Matsushima & Graybiel, 2020). sSPNs and mSPNs exhibited marked differences in their encoding of outcomes. Outcome encoding by sSPNs was particularly strong when RPE or PPE was high, a finding in accord with the proposed role for striosomes as being part of the circuit that calculates the learning signals (Doya, 2000). Notably, we did not find differential encoding of motor behavior. This finding suggests that the learning function may be a principal function of striosomes added to other shared attributes with neurons of the surrounding matrix (Amemori et al., 2011; Friedman et al., 2020).

We suggest that multiplex encoding of action-outcome associations and PEs are key characteristics of large numbers of SPNs in the anterodorsal striatum, that striosomes in this region are particularly sensitive to error signals of both positive or negative valence outcomes, and that models incorporating these features could be of great value in understanding how striatal circuits underpin adaptive behaviors in uncertain environments.

## RESULTS

### A New Dynamic Foraging Task with Rewarding and Aversive Outcomes

Mice were trained on the CBB task with rapidly changing action-reward and action-air puff contingencies. Head-fixed mice, with their forepaws on a wheel (Figure 1A), initiated trials by holding the wheel still for 2 s (Figure 1B). Mice had 3 s to make a leftward or rightward response. Both actions were probabilistically linked to receipt of a rewarding outcome (water) and/or to the receipt of punishment (air puff delivered to the face). The action-reward and action-puff contingencies changed rapidly without cueing, and the reward and puff block changes were made independent of one another (Figure 1C) after mice received 6-15 rewards or avoided this number of puffs. Reward and puff action-outcome contingencies switched independently of one another; actions could lead to one of four outcome pairs (reward-puff, reward-no puff, no reward-puff, and no reward-no puff). Mice performed hundreds of trials (average: 366; range: 247-585) in each session, with tens of reward (mean: 20.4; range: 14-34) and puff (mean: 22.9; range: 14-38) block switches. They reliably adapted their behavior within blocks with a given action-reward or action-puff contingency (Figure 1D) and around action-reward or action-puff contingency switches (Figure 1E). We compared choice behavior as a function of the actions and outcomes of the previous two trials (Figure 1F), and we performed an autoregression analysis using the last five trials (Figure 1G). Both analyses show that mice incorporate outcomes of multiple past trials rather than relying on a strict win-stay/lose-shift strategy.

**Figure 1.**
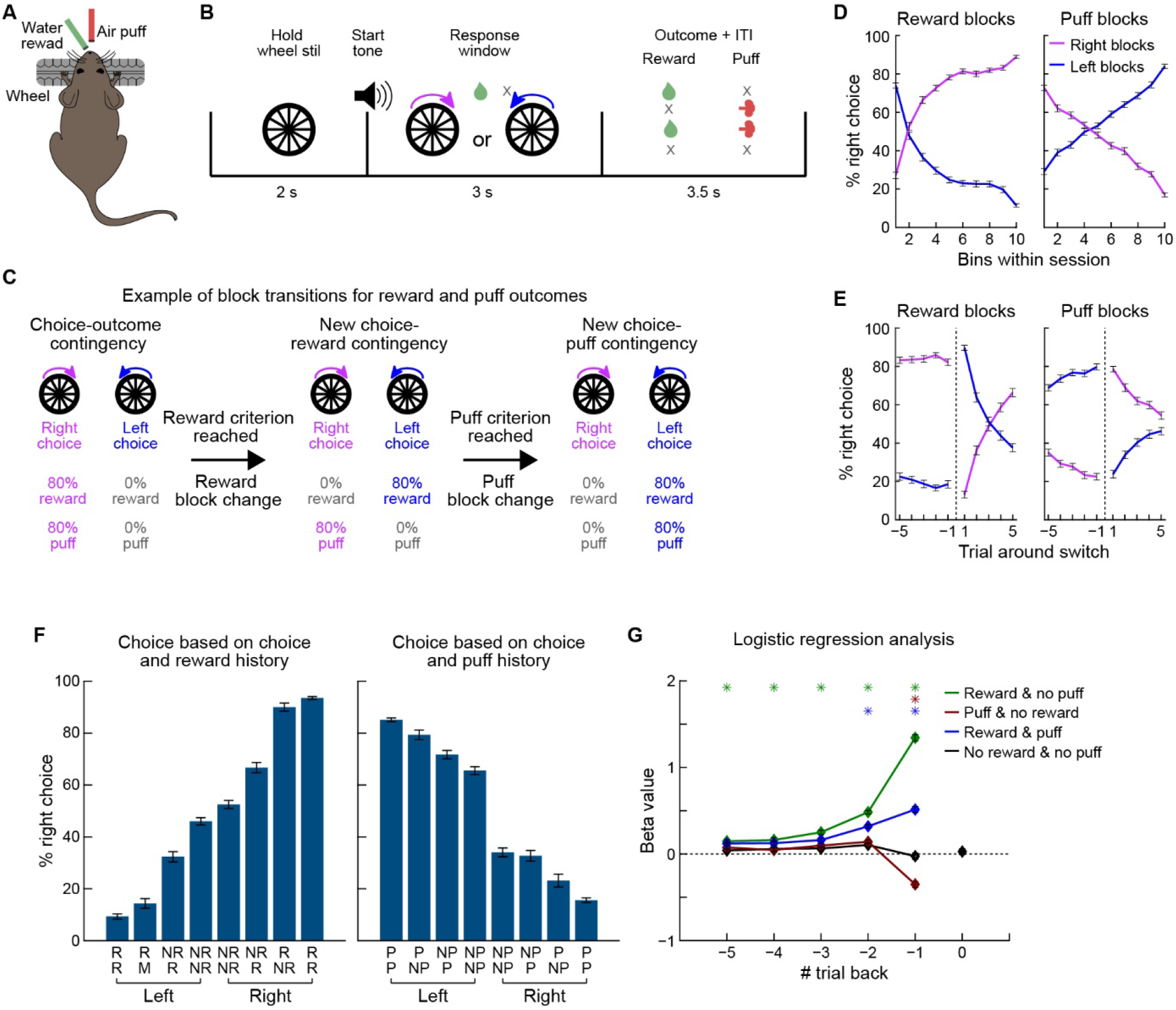
The Cost-Benefit Bandit Task. (A) Behavioral setup. Head-fixed mice reported choices by rotating a wheel to the left or the right. Water rewards and air puffs were delivered via two tubes. (B) Trial procedure. Mice initiated trials by holding the wheel still for 2 s. An auditory go-cue signaled trial start, after which the mouse had 3 s to move the wheel 15° in either direction. Left and right actions were probabilistically linked to reward and air puff delivery, giving four possible outcome combinations: reward-no puff, puff-no reward, reward-puff, and no reward-no puff. (C) Action-reward and action-puff contingencies changed in block of trials independently of each other. (D) Choices within the reward and puff blocks. Trials within blocks were grouped into 10 bins. Error bars show SEM across sessions (n = 75 sessions from 13 mice). (E) Behavioral adaption during five trials before and after reward and puff block switches. (F) Mouse decisions were affected by the outcomes and actions in the last two trials. For simplicity, only trials in which the mouse made the same action in the last two trials were included (R, reward; NR, no reward; P, puff; NP, no puff). (G) Regression coefficients (mean ± SEM) for the 4 types of outcomes ( reward/no puff, puff/no reward, reward/puff, and no reward/no puff) for the preceding five trials, as calculated using an auto-regressive model. n = 13 mice. *p < 0.05, t-test comparing the beta values of reward, puff and both outcomes with no outcome.

### Single Striatal Neurons Encode Associations between Actions and Multiple Conflicting Outcomes

We used 2-photon imaging to measure Ca^++^ activity of GCaMP6s-expressing SPNs via a cannula placed above the striatum (Figure 2A). In every session, we recorded a new field of view, resulting in a total of 5831 unique neurons (75 sessions in 13 mice). Every field of view contained SPNs of both striosome (n = 2249) and matrix (n = 3582) compartments (Figure 2B). Only low percents of striosomal neurons were labeled, but with this preparation we could identify the compartments based on labeling of the neuropil. We regarded every neuron in a red-labeled neuropil cluster as being a striosomal SPN. To test for behavioral effects of implanting the imaging cannula, we compared the performance of mice used for imaging to mice without an implanted imaging cannula. There was no significant effect of the cannula on the number of sessions required for training the CBB task (Figure S1A). In trained mice, the response time was significantly higher in mice with a cannula, but otherwise the performance of the mice was identical (Figures S1B-S1E).

**Figure 2.**
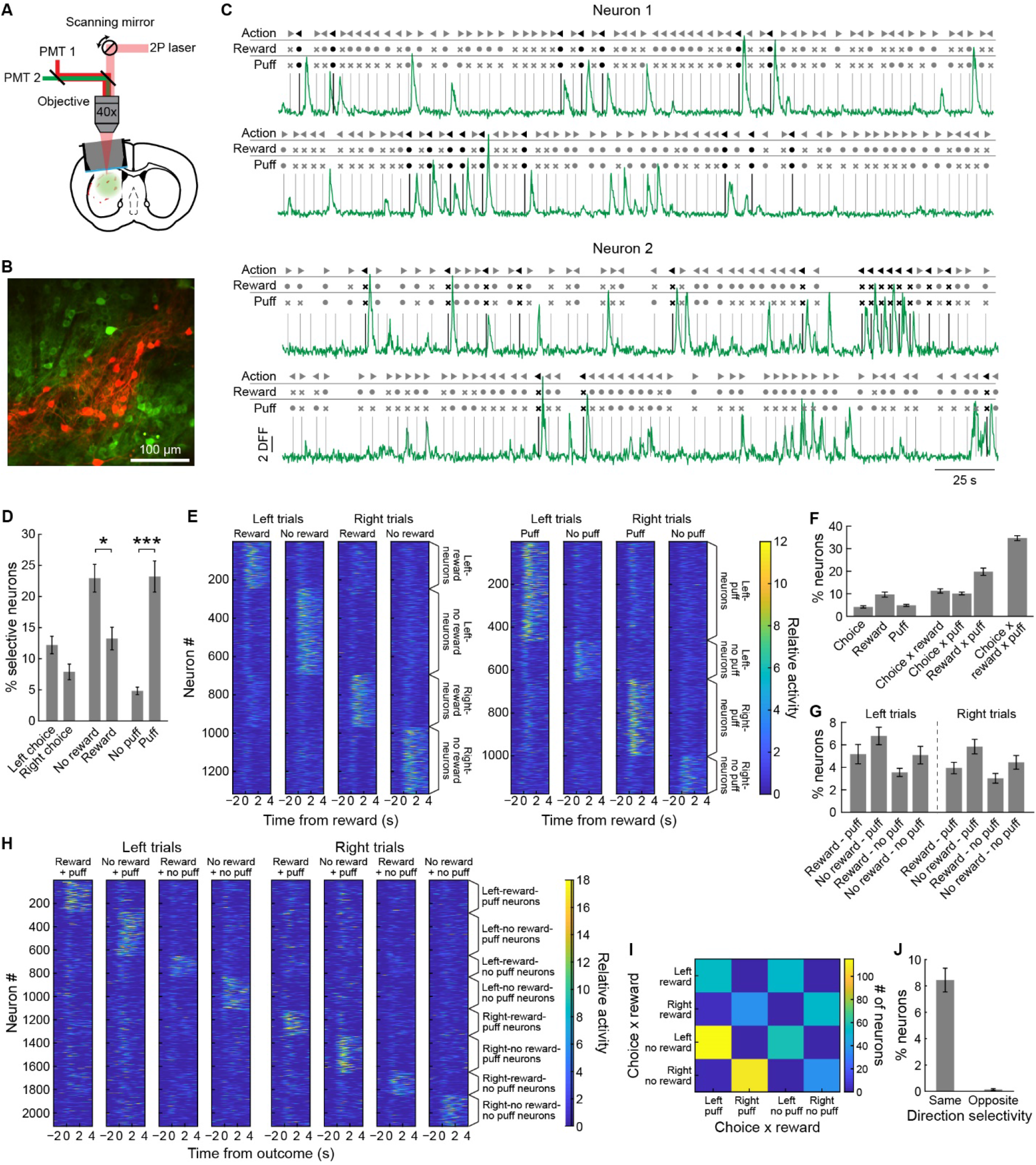
Representations of Action-Reward-Puff Combinations by SPNs. (A) Imaging setup and preparation. (B) Example of imaging field-of-view showing GCaMP6s-expressing neurons (green). tdTomato (red) expressed in the cell bodies and neuropil demarcates the striosome compartment. (C) Two sample neurons during 10 min of recording. Neuronal transients occurred selectively for action-reward-puff associations during the outcome period. (D) Percentage (mean ± SEM) of neurons selectively responding to chosen action, reward outcomes, and puff outcomes. n = 13 mice for all panels. *p < 0.05; ***p < 0.001; paired t-test. (E) Activity of neurons that represented action-outcome associations, averaged over all trials for a given action-outcome combination per session. Activity of each neuron was normalized to its average transient rate for all trials (left: action-reward representations; right: action-puff representations). (F) Percentage of neurons per mouse with chosen action, reward, puff, 2-way interactions, and 3-way interactions as significant predictors in a stepwise regression analysis. (G) Percentage of neurons with activity selective for the 8 possible action – reward outcome – puff outcome combinations, identified in a stepwise regression analysis. (H) Average activity of the neurons in D, shown separately for the 8 different trial types. Neuronal activity was normalized to the average transient rate. (I) Joint distribution of the number of neurons responding to different action-reward and action-puff associations. (J) Among the neurons that selectively responded to both an action-reward and action-puff associations, the vast majority had a stable action-preference (p < 0.001, paired t-test).

We identified individual Ca^++^ events in ΔF/F (DFF) traces using a custom algorithm (Methods, Figure S2A). Using a chi-square test, we compared the number of trials in which a Ca^++^ transient was observed in the 3 s following outcome delivery to identify neurons with selective responses to specific actions and outcomes. Many striatal neurons showed strong activity that was selective for rewarding outcomes (reward: 13.1 ± 1.8%, no reward: 22.7 ± 2.2%), puff outcomes (puff: 23.0 ± 2.5%, no puff: 4.8 ± 0.6%) and chosen actions (left: 12.1 ± 1.4%, right: 7.8 ± 1.2%; mean percentage of neurons per mouse averaged over 13 mice; Figures 2C and 2D).

Following evidence that SPNs encode action-outcome associations for reward (Lau & Glimcher, 2007; Stalnaker et al., 2010), we tested whether reward- and puff-outcome activity was selective for specific actions. We took two complementary approaches. First, for every neuron, we used a stepwise logistic regression analysis to determine which factor(s) among the chosen action and received outcomes could best predict whether or not a transient occurred in a trial (using a cutoff of p < 0.05). We then identified neurons encoding a factor as neurons that included that factor in their model. The activity of the neurons that were identified in this analysis, averaged over all trials for a given action-outcome combination, is shown in Figure 2E. In addition, we used a more conservative approach, employing sequential chi-square analyses. With this approach, we found many reward-selective and puff-selective SPNs among the action-selective SPNs (left: reward: 14.9 ± 2.9%, no reward: 16.8 ± 1.6%, puff: 16.6 ± 2.7%, no puff: 6.6 ± 1.9%; right: reward: 17.2 ± 3.4%, no reward: 25.0 ± 3.9%, puff: 30.9 ± 4.8%, no puff: 9.7 ± 2.6%; mean percentage of neurons per mouse averaged across 13 mice; Figures S2D and S2E) and action selective neurons among reward- and puff-selective neurons (Figures S2F and S2G). Finally, we used a regression analysis to find the regression coefficients for actions, reward and puff outcomes for every neuron imaged. (Figures S2H-S2I). These findings indicate that the striatal neurons encoded action-puff contingencies similarly to their encoding of action-reward contingencies.

Next, we asked whether action-reward and action-puff contingencies were represented by separate or overlapping populations of SPNs. We found in our stepwise regression analysis that 34.66 ± 1.1% of neurons (mean ± SEM, n = 13 mice) encoded combinations of actions and both rewards and puffs (Figure 2F). We used a second regression analysis to identify the neurons that encoded each of the 8 possible action-reward-puff associations (Figure 2G, S2J). The average activity of action-reward-puff neurons for the 8 possible trial types (chosen action x reward outcome x puff outcome) showed selective responses for specific combinations (Figure 2H). We noted that neurons that encoded all 3 factors had stable action encoding. Among the neurons that encoded both an action-reward and action-puff association, almost all encoded both outcomes for the same action (Figures 2I and 2J; same action: 8.5 ± 0.9% of all neurons, different: 0.1 ± 0.1%; n = 13 mice, p < 0.001, paired t-test). These analyses demonstrated that SPNs can encode associations of multiple outcomes with a given action.

### Multiplexed Encoding of Reward and Aversive Outcomes of Actions in Striatal Neurons

The responses of the individual SPNs could represent outcome value (i.e., value coding) if the activity signaled good (reward and/or no puff) or bad outcomes (no reward and/or puff). Alternatively, action-outcome encoding could be multiplexed such that, as a population, the SPNs would respond to all combinations of actions and outcomes without being biased towards specific combinations. In this case, partially overlapping populations of single SPNs would encode each action-outcome contingency, and single SPNs encoding both outcomes would not necessarily represent the combined value of the outcomes. Our results favor this latter form of multiplexed action-outcome encoding (Figure 3).

**Figure 3.**
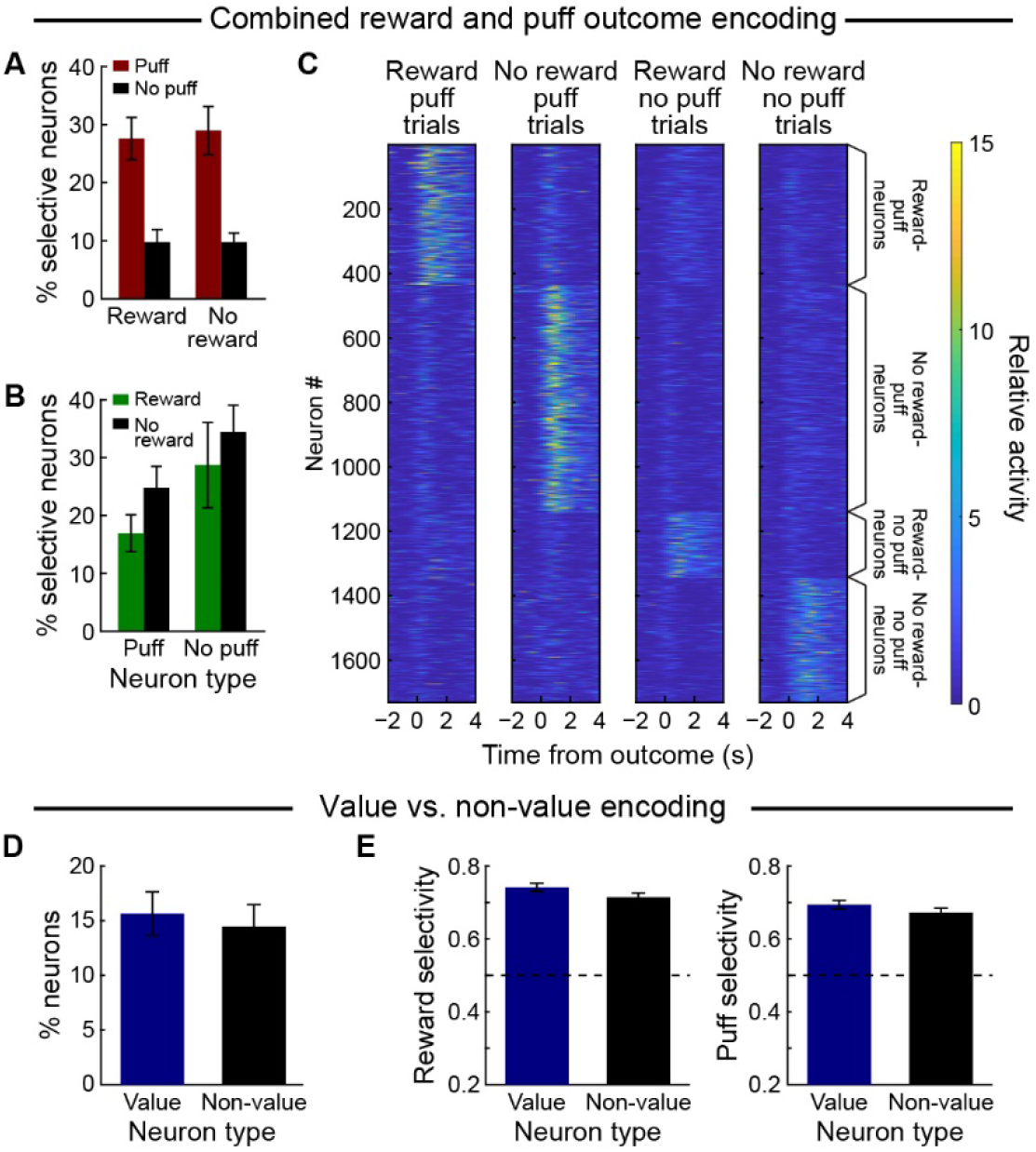
Combined Reward-Puff Representations in Single Neurons Do Not Always Reflect Outcome Value. (A) Percentage (mean ± SEM) of reward- and no-reward-selective neurons that were selective for the puff outcome, as identified using chi-square analysis, (n = 13 mice for all panels). There was a significant main effect of the percentage of puff neurons (p < 0.001, repeated measures ANOVA) but no reward-puff interaction (p = 0.83). (B) Percentage of puff- and no-puff-selective neurons that were selective for reward/no-reward outcome. There was a significant main effect of puff neurons (p < 0.05, repeated measures ANOVA) but no interaction between reward and puff (p = 0.84). (C) A stepwise regression analysis identified neurons with reward and puff outcome interactions. Session-averaged activity of these neurons is shown for trials with the different reward – puff combinations. (D) Neurons were split into value and non-value type based on whether they responded to reward and puff oppositely or not. The proportion of neurons classified to these two types was not different. (E) The selectivity of value and non-value neurons in D is not significantly different for reward (left) or puff (right).

Many single SPNs responded in relation to both reward and puff or to the absence of both. Chi-square analysis showed that the percentages of puff-selective and no-puff selective SPNs were not different among the reward neurons (puff: 27.6 ± 3.6%; no puff: 9.7 ± 2.2%, average percentage per mouse) and no-reward neurons (puff: 29.0 ± 4.2%; no puff: 9.7 ± 1.6%; Figure 3A). Conversely, the distribution of reward and no-reward selective SPNs was similar among the puff- (reward: 16.9 ± 3.2%; no reward: 24.8 ± 3.7%) and no-puff-selective (reward: 28.7 ± 7.4%; no reward: 34.5 ± 4.6%) neurons (Figure 3B). With ANOVA, we found no interaction in the distribution of puff/no-puff neurons amongst the reward/no-reward population (Figure 3A; p = 0.83, n = 13 mice) or vice versa (Figure 3B; p = 0.84). These results suggest that the selectivity of a given SPN for one outcome did not depend on the selectivity for the other outcome. The activity of all neurons responding to both outcomes is shown in Figure 3C, averaged over all trials for each of the 4 different reward-puff combinations.

To compare value encoding and multiplexed outcome encoding directly, we identified ‘value neurons’, i.e., SPNs whose activity reflected that an action was good or was bad (good outcomes: reward and no puff; or bad outcomes: puff and no reward), and ‘non-value neurons’, i.e., SPNs that responded in relation to the presence or absence of both outcomes. We found similar proportions of value and non-value neurons among the recorded neurons (value: 15.6 ± 2%, non-value: 14.5 ± 2%, average percentage across 13 mice; p > 0.05, paired t-test, Figure 3D). The value and non-value neurons did not detectably differ in their selectivity for reward (value: 0.74 ± 0.01, non-value: 0.71 ± 0.01, p > 0.05) or puff (value: 0.69 ± 0.01, non-value: 0.67 ± 0.01, p > 0.05; Figures 3E and S3A).

The representation of both ‘good’ and ‘bad’ outcomes in single neurons could potentially be accounted for by difference in action selectivity for both outcomes. For example, a neuron could be active in trials where left choices lead to reward or right choices lead to air puffs. This possibility appeared unlikely, because we found stable action selectivity (that is, the same direction of movement selectivity for both action-outcome associations) (Figures 2J). We further tested this idea, however, by quantifying the activation of neurons encoding action-reward-puff interactions in all eight action-reward-puff trial types. SPNs that encoded the good or bad outcome of one action, as a population, were not activated when the mouse selected the opposite action and received outcomes of opposite value (Figure S3B). Hence, the population activity of the striatal SPNs encoded combinations of outcomes in a multiplexed manner so that their activity reflected multiple outcomes of actions rather than the overall value of an action.

### Encoding of Multiplexed Prediction Errors Derived from a Cost-Benefit Reinforcement Learning Model

We used RL models to gain more insight into the behavior and to test how trial-by-trial PEs for rewarding and aversive outcomes were represented in the SPNs. In such models, costs and rewards associated with actions are typically combined into one scalar value, and for every action one action-value is learned. However, the observed activity of SPNs in mice performing the CBB task suggested that these outcomes could be represented in parallel for cost-based and reward-based associations. We therefore adapted existing RL models to integrate costs and benefits using two alternative approaches (see Methods). In the first model, one set of Q-values is used to model the expected overall value of the two possible actions. When outcomes are delivered, reward and puff outcomes are weighted and combined into one outcome value, which is used to form one PE, and to update one set of action values. We refer to this model as the *integrated model*. In the second model (Figure 4), two parallel sets of Q-values are estimated, one for reward and one for puff. PEs are calculated separately for both outcomes, and the two sets of Q-values are updated in parallel using two sets of learning rates, for reward and for puff. We refer to this model as the *parallel model*.

**Figure 4:**
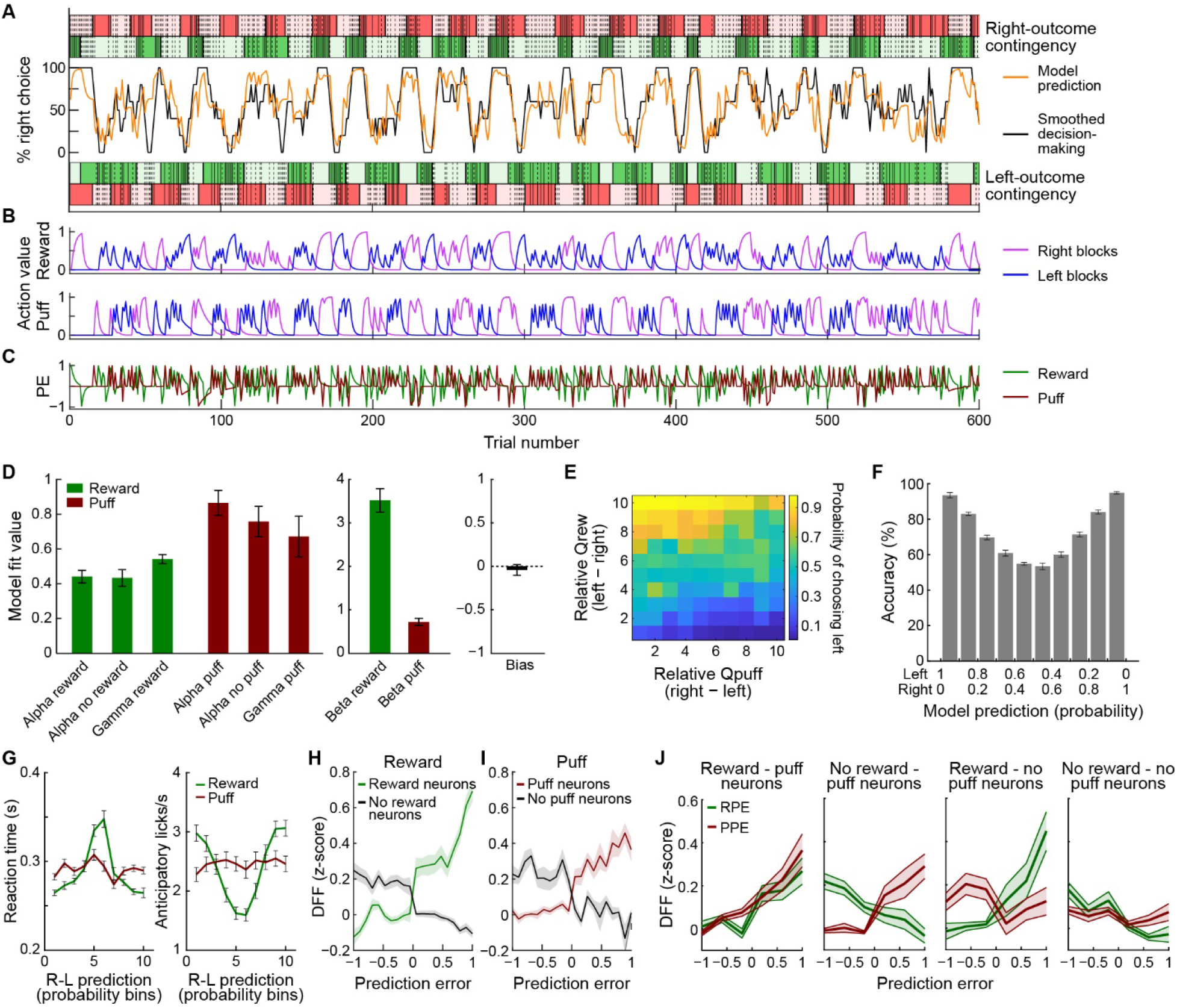
Outcome Neurons Are Modulated by RPE and PPE. (A) Session example. Action-outcome contingency is shown as colored blocks (reward: green; puff: red) for right (top) and left (bottom) choices. Dark and light red/green shading indicates, respectively, 80% and 0% probability of outcome delivery. Within blocks, solid lines indicate trials in which the outcome was delivered, and dotted lines indicate trials in which the outcome was not delivered. Middle lines show the smoothed decisions by mice (black) and model prediction (orange). (B) Estimated Q-values for the reward (top) and puff (bottom) for both actions for every trial. (C) RPE and PPE calculated for every trial by subtracting the Q-value of the chosen action from the outcome. (D) Average model parameters estimated by the RL model (alpha: learning rate for trials with reward/no reward and puff/ no puff; gamma: outcome specific forgetting rate; beta: inverse temperature). (E) Fraction of trials in which the mouse chose the left action as a function of relative reward Q-values and relative puff Q-values. (F) Accuracy as a function of the probability of left and right actions as predicted by the model. (G) Trial-by-trial model predictions correlated with reaction time (left) and anticipatory licking (right). (H) ΔF/F (DFF) response during outcome period was correlated with RPE in reward neurons (green, r = 0.89; p < 0.001) and in no-reward neurons (black, r = 0.76; p < 0.05). (I) DFF was significantly correlated with PPE in puff neurons (red, r = 0.92; p < 0.001), but not in no-puff neurons (black, n.s.). (J) Activity of neurons that responded to both reward and puff outcomes (Figure 3C) is modulated by RPE and PPE (reward – puff neurons: RPE: r = 0.68, p < 0.01; PPE: r = 0.68, p < 0.01; no-reward – puff neurons: RPE: r = −0.82, p < 0.001; PPE: r = 0.45, p < 0.05; reward – no-puff neurons: RPE: r = 0.88, p < 0.001; PPE: r = −0.11, p < 0.05; no-reward – no-puff neurons: RPE: r = −0.40, p < 0.05; PPE: r = −0.5; p = 0.05).

The parallel model performed slightly but non-significantly better, as shown using cross-validation (Figure S4A). We tried reducing the number of parameters by having the same learning rates for reward and puff, or by reducing the number of separate learning rates for reward or puff, but both reduced the accuracy of the model in a test set (Figure S4B). We also tested the history dependency of decision-making by setting the learning rates to 1, resulting in a win-stay/lose-shift model, and again found that this significantly impaired model performance.

Next we tested whether the PE variables from the two models could account for the neuronal activity using a stepwise logistic regression model. We found that reward and puff outcomes could account for firing in more neurons than could the integrated value of the outcome (Figure S4C). Similarly, RPE and PPE were predictive of firing in more neurons than an integrated PE. Therefore, we used the parallel cost-benefit RL model to derive reward- and puff-specific action values and prediction errors (Figures 4A-4D).

Using this model, we found that the relative difference in the inferred positive and negative action values predicted the selected actions of the mice (Figures 4E and 4F). The model predictions were systematically correlated with reaction times and anticipatory licking in the 1-s period preceding the outcome (Figure 4G). Hence, the model predictions could be generalized to behavioral variables beyond the choices and their outcomes that were used to fit the model.

We then further characterized the modulation of the observed reward and puff outcome activity by RPE and/or PPE. The activity of reward-responsive and puff-responsive neurons was correlated with model-derived RPE (reward neurons: r = 0.89, p < 0.001; no-reward neurons: r = −0.76, p < 0.05; Figure 4H) and PPE (puff neurons: r = 0.92, p < 0.001; no-puff neurons: r = −0.51, p > 0.05; Figure 4I) during the 3-s window after outcome delivery. In neurons modulated by both reward and puff outcomes, as found using stepwise regression analysis (Figure 2), outcome-related activity was modulated by both RPE and PPE (reward – puff neurons, RPE: r = 0.68, p < 0.01; PPE: r = 0.68, p < 0.01; no-reward – puff neurons, RPE: r = −0.82; p < 0.001; PPE: r = 0.45; p < 0.05; reward – no-puff neurons, RPE: r = 0.88; p < 0.001; PPE: r = −0.11; p < 0.05; no-reward – no-puff neurons: RPE: r = −0.40; p > 0.05; PPE: r = −0.5; p < 0.05; Figure 4J). The modulation of neuronal activity was observed in the period after the outcome was delivered. However, instead of reflecting prediction errors, this activity could be produced by differences in reward and puff anticipation before outcome delivery because of the slow dynamics of Ca^++^ signals. We consider this possibility unlikely, as if so, we would have expected higher neuronal activity in trials with high reward/puff expectation, which is the opposite of what was observed.

### Stronger Encoding of Reward and Puff Outcomes in Striosomes Than in the Surrounding Matrix

We tested the hypothesis that mSPNs differentially encode ‘motor’ and and sSPNs ‘limbic’ reinforcement-related information. First, we identified action- and outcome-encoding neurons using chi-square analysis (Figure 5A). More sSPNs than mSPNs were selectively activated for reward (sSPNs: 14.7 ± 1.4%, mSPNs: 11.7 ± 1.9%, p < 0.05, repeated measures t-test based on the average per mouse), or puff (sSPNs: 24.9 ± 2.2%, mSPNs: 22.0 ± 2.8%, p < 0.05) or no-puff (sSPNs: 6.1 ± 0.7%, mSPNs: 4.2 ± 0.6%, p < 0.05) outcomes. However, we did not find a compartmental difference between the number of neurons encoding chosen actions (left: sSPNs: 11.7 ± 1.1%, mSPNs: 12.7 ± 1.7%; right: sSPNs: 9.1 ± 1.4%, mSPNs: 7.5 ± 1.3%, t-test on averages per mouse). The distribution of single-neuron regression coefficients concurred with these results (Figure 5B); we observed significant differences in outcome encoding between the striosomal and matrix populations (reward: p < 0.005, puff: p < 0.01; n = 2249 sSPNs and 3582 mSPNs, Kolmogorov–Smirnov test), but not in action encoding.

**Figure 5.**
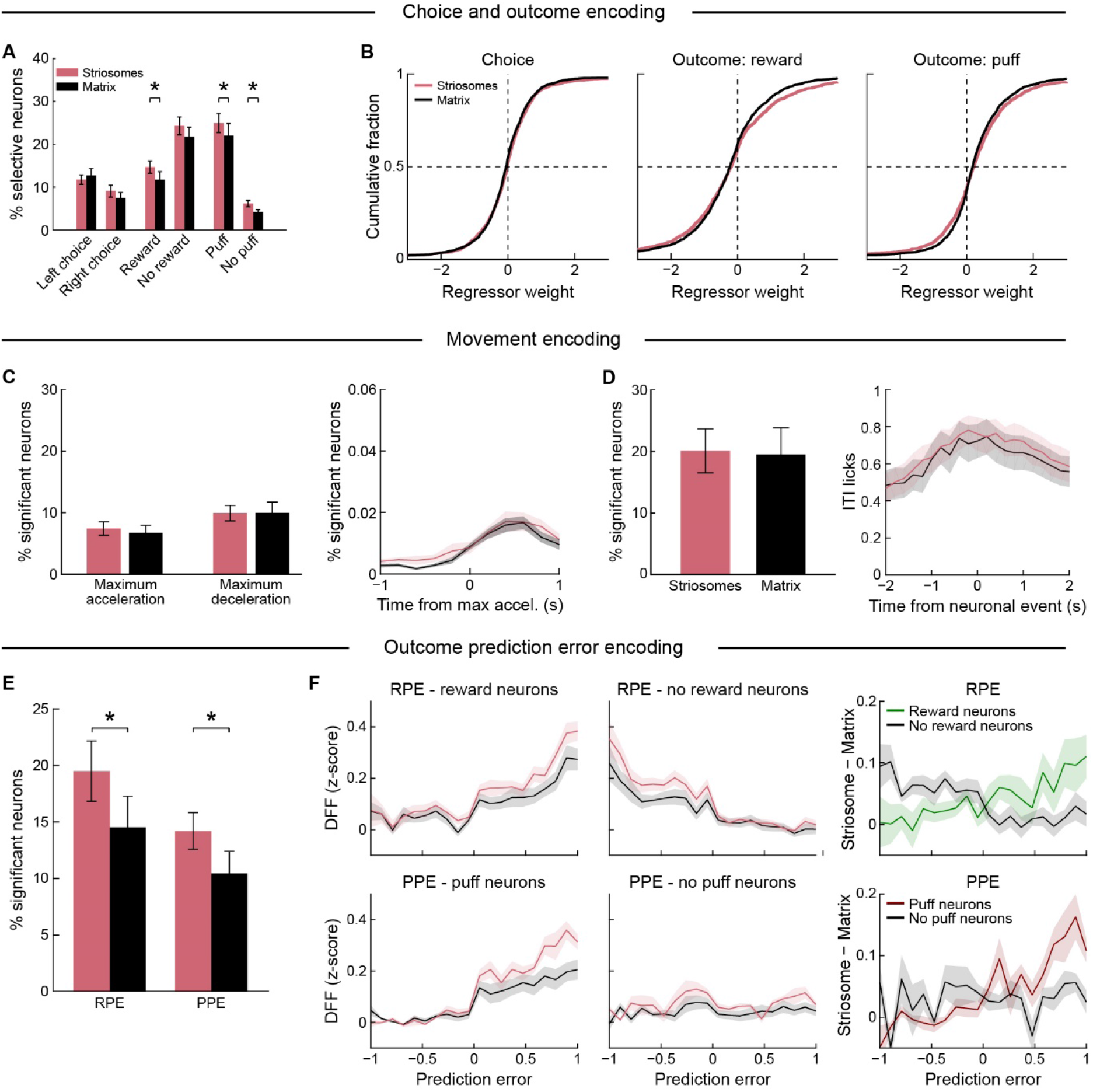
Preferential Outcome and Prediction Error Representation for Rewards and Punishment in Striosomes. (A) Percentage (mean ± SEM) of neurons selective for chosen action, reward or puff outcome (n = 13 mice for all panels). More sSPNs were activated by reward outcomes, puff and no-puff outcomes (*p < 0.05, paired t-test). There was a trend for no-reward neurons (p = 0.07). (B) Regression coefficients of sSPNs (n = 2249) and mSPNs (n = 3582). The distribution of sSPNs and mSPNs was significantly different for reward (p < 0.005) and puff outcomes (p < 0.01) but not for chosen action (Kolmogorov-Smirnov test). (C) Movement modulation was assessed by aligning neuronal activity to maximum acceleration and decelaration of the wheel during ITIs. Percentage of sSPNs (red) and mSPNs (black) with significant movement modulation (left), and average responses of all modulated neurons, aligned to the time of maximum acceleration (right). (D) Licking during ITIs was aligned to neuronal activity (detected transients). Panels show percentage of sSPNs and mSPNs with significantly more activity-triggered licking than expected by chance (left), and licking aligned to neuronal activity for significantly modulated neurons (right). (E) Average percentage of sSPNs and mSPNs per mouse that were modulated positively by RPE or PPE in the stepwise regression model. *p < 0.05. (F) RPE and PPE modulation of DFF in sSPNs and mSPNs that were selective for reward outcome (top) or puff (bottom) outcome, respectively. The correlation coefficient for sSPNs was significantly higher than for mSPNs (reward: p < 0.05; puff: p < 0.01, n = 13 mice). Right: the difference between the modulation in the sSPNs and mSPNs was significantly correlated with RPE in reward neurons (r = 0.77; p < 0.01, n = 13 mice) and PPE in puff neurons (r = 0.72; p < 0.05, n = 13 mice).

Task-related actions are strongly linked to outcomes. We therefore tested whether mSPNs had a stronger encoding of movements than sSPNs during intertrial intervals (ITIs), times during which the actions were likely to be less related to task performance proper than during the trials themselves. We identified the onset and offset of wheel movement bouts during ITIs. The similar proportions of neurons in each compartment were modulated by movement onset (sSPNs: 7.4 ± 1.1%, mSPNs: 6.7 ± 1.2%, t-test, n = 13 mice) and offset (sSPNs: 9.9 ± 1.3%, mSPNs: 10.0 ± 1.8%) regardless of movement direction (Figures S5A-S5F). Moreover, similar proportions of sSPNs and mSPNs were modulated for maximum acceleration or deceleration of wheel movements during ITIs (Figure 5C). We similarly aligned neuronal activity to the onset of licking bouts during the entire session (sSPNs: 22.1 ± 3.6%, mSPNs: 22.0 ± 3.0%) or ITI period (sSPNs: 4.7 ± 0.9, mSPNs: 3.7 ± 1.3) and found no differences in the proportion of responsive sSPNs and mSPNs (Figures S5G and S5H). Finally, we aligned licks during ITIs to the time of neuronal events to quantify the number of neurons with activity coincident with licking (Figure 5D) and again found no difference (sSPNs: 4.7 ± 1.3%, mSPNs: 3.7 ± 0.9%). These results point to a similar activation of sSPNs and mSPNs in relation to movements during trials and ITIs.

Striosomes have been proposed to function as critics in actor-critic RL models (Doya, 2000). Our finding that sSPNs had a stronger encoding of both positive and negative outcomes is consistent with this view. For a further test of this proposition, we asked how SPN activity in each compartment was modulated by RPE and PPE. We used regression analysis to identify neurons for which activity could be better accounted for if RPE and/or PPE were included as regressors. The percentage of sSPNs was higher for both RPE (sSPNs: 19.5 ± 2.7%, mSPNs: 14.5 ± 2.8%, p < 0.05, repeated measures t-test on n = 13 mice) and PPE (sSPNs: 14.2 ± 1.6, mSPNs: 10.4 ± 1.9%, p < 0.05; Figures 5E and S5I). Moreover, reward- and puff-responsive neurons in striosomes were more strongly modulated by, respectively, RPE and PPE than those in the matrix (Figure 5F). As a result, the relative sSPN-to-mSPN activation was correlated with RPE (reward neurons: r = 0.77, p < 0.01; no-reward neurons: r = −0.62, p = 0.06) and PPE (puff neurons: r = 0.72, p < 0.05; no-puff neurons: r = 0.3, p = n.s.). This set of results is compatible with a notion of striosomes as having sufficient information about RPE and PPE to provide teaching signals for downstream circuits.

We also tested whether there were differences in the proportion of sSPNs and mSPNs with activity related to action-outcome associations. By regression analysis, we did not find a difference (Figures S5J-S5L). This result suggests that SPNs in both compartments can represent associations between actions and outcomes. Thus, our findings indicate that outcome and prediction errors signaling is stronger in sSPNs, but that responses related to movement per se can be similarly detected in each population.

### Action Outcomes Can Be More Reliably Decoded Using the Activity of sSPNs Than by the Activity of mSPNs

The single-cell SPN data indicated that sSPNs exhibited preferential outcome-related activity, but that there was no clear compartmental difference in encoding of motor behavior. Yet it was still possible that differences between striosome and matrix compartments in movement-related activity would emerge when analyzing population activity. We therefore used decoding analyses to evaluate which task-relevant information could be read out by downstream structures from the striatal population activity. We trained artificial neural networks (ANNs) to classify trials with each of the eight possible action-outcome combinations. We trained separate models using all SPNs or only sSPNs or mSPNs (Figure 6A). We first performed a decoding analysis using pseudo trials, constructed by concatenating the activity of SPNs recorded across all sessions (n = 75 sessions). Pseudo-trial-based models predicted the action-reward-puff combination with very high accuracy (all SPNs: 96%; sSPN: 93%; mSPN: 90%; Figure 6B).

**Figure 6.**
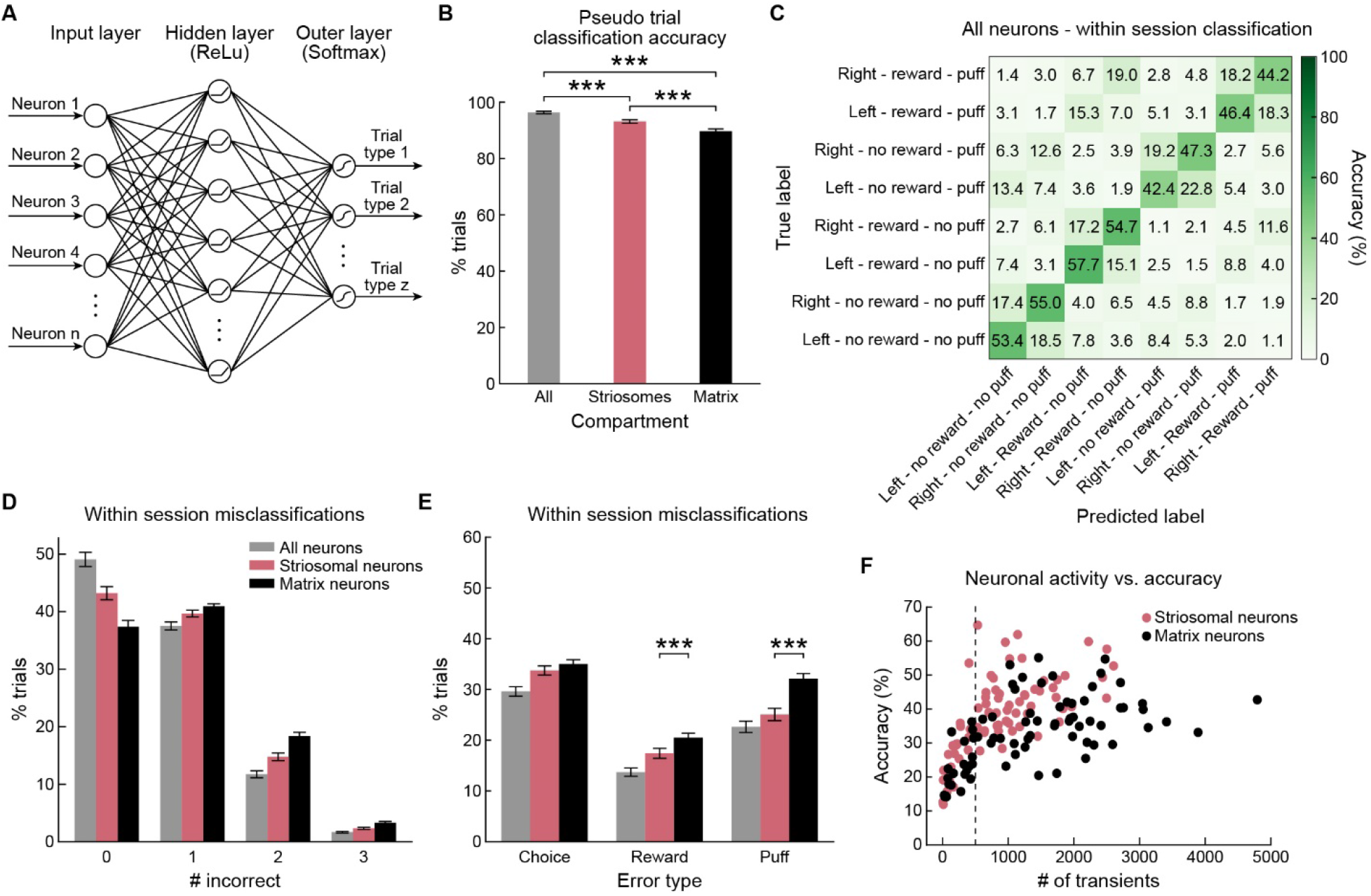
Decoding Analysis of Action-Outcome Combinations. (A) Artificial neural network architecture used for behavioral decoding. The hidden layer had the same size as the input layer. (B) Accuracy (mean ± SEM) of model using pseudo-trials for all SPNs (n = 5831) and subsampled sSPNs (n =2249) or mSPNs (n =2249). The model including all SPNs had a higher accuracy than the compartment-specific models (***p < 0.001), and the striosome model outperformed the matrix model (p < 0.001). (C) Confusion matrix of session-based models showing percentage of trials with the true and predicted label for each of the 8 different trial types. (D) Percentage of trials in which 0, 1, 2 or 3 dimensions were incorrectly predicted by session-based models. (E) The models based on sSPN activity significantly outperformed the models based on mSPN activity when predicting reward and puff outcome but not chosen action (***p < 0.001). The model based on all neurons had an accuracy higher than the matrix model for all three features (p < 0.001) and higher than the striosome model for chosen action (p < 0.005) and reward (p < 0.01), but the performance was not significantly different for puff. (F) Relationship between the model accuracy and the total number of transients recorded across all imaged neurons per session. Some sessions had a lower number of transients, due to low activity or a low number of neurons. Sessions on the left of the dashed line were excluded from the decoding analysis.

Because pseudo trials can inflate decoding accuracy by decoupling behavioral and neuronal variability, we also trained ANNs for individual sessions. We could accurately decode the action-outcome combination in 49% of the trials (chance level: 12.5%) using the activity of all SPNs within a session (n = 77.75 ± 4.16 neurons, 75 sessions; Figure 6C), and 37% and 43%, respectively, using matching numbers of sSPNs and mSPNs (n = 28.68 ± 1.82 neurons, 75 sessions; Figures S6A and S6B). We quantified the percentage of trials in which the chosen action, the reward outcome or the puff outcome was misclassified (Figures 6C-6E). As expected, mSPN-based models had significantly more misclassifications for the reward (sSPNs: 17.4 ± 1.0%; mSPNs: 20.5 ± 0.9, p < 0.05) and puff outcome (sSPNs: 25.1 ± 1.2%; mSPNs: 32.1 ± 1.0%, p < 0.05) than sSPN-based models, but there was no difference for chosen action (sSPNs: 33.7 ± 0.9 and mSPNs: 35.0 ± 0.9%, Figure 6E). The higher accuracy for sSPNs was observed regardless of the total number of transients that were observed in a session, indicating that differences in transient rates cannot account for the differential accuracy (Figure 6F).

These decoding analyses demonstrate that SPN ensemble activity contains robust information about action-outcome contingencies, and they confirm the conclusion based on our previous analyses (Figure 5) that the sSPNs that we imaged exhibited stronger activity in relation to the outcome of actions than did the mSPNs imaged simultaneously in the same fields of view.

### Decoding Future Actions Using SPN Activity

Updating of action values allows agents to adapt their behavior in order to maximize value. We tested whether the recorded activity contained information about future decision-making. Some SPNs responded differentially depending on whether the mouse was going to stay with its action or switch in the next trial (Figure 7A). More SPNs encoded future switching than stay behavior during the outcome period (sSPNs and mSPNs,chi-square analysis, p < 0.001; Figure 7B). In agreement with the hypothesis that striosomes are important for cost-benefit decision-making (Friedman et al., 2015, 2017), we found significantly more sSPNs than mSPNs encoding future switching selectively in trials with a combined reward and puff outcome (p < 0.05).

**Figure 7.**
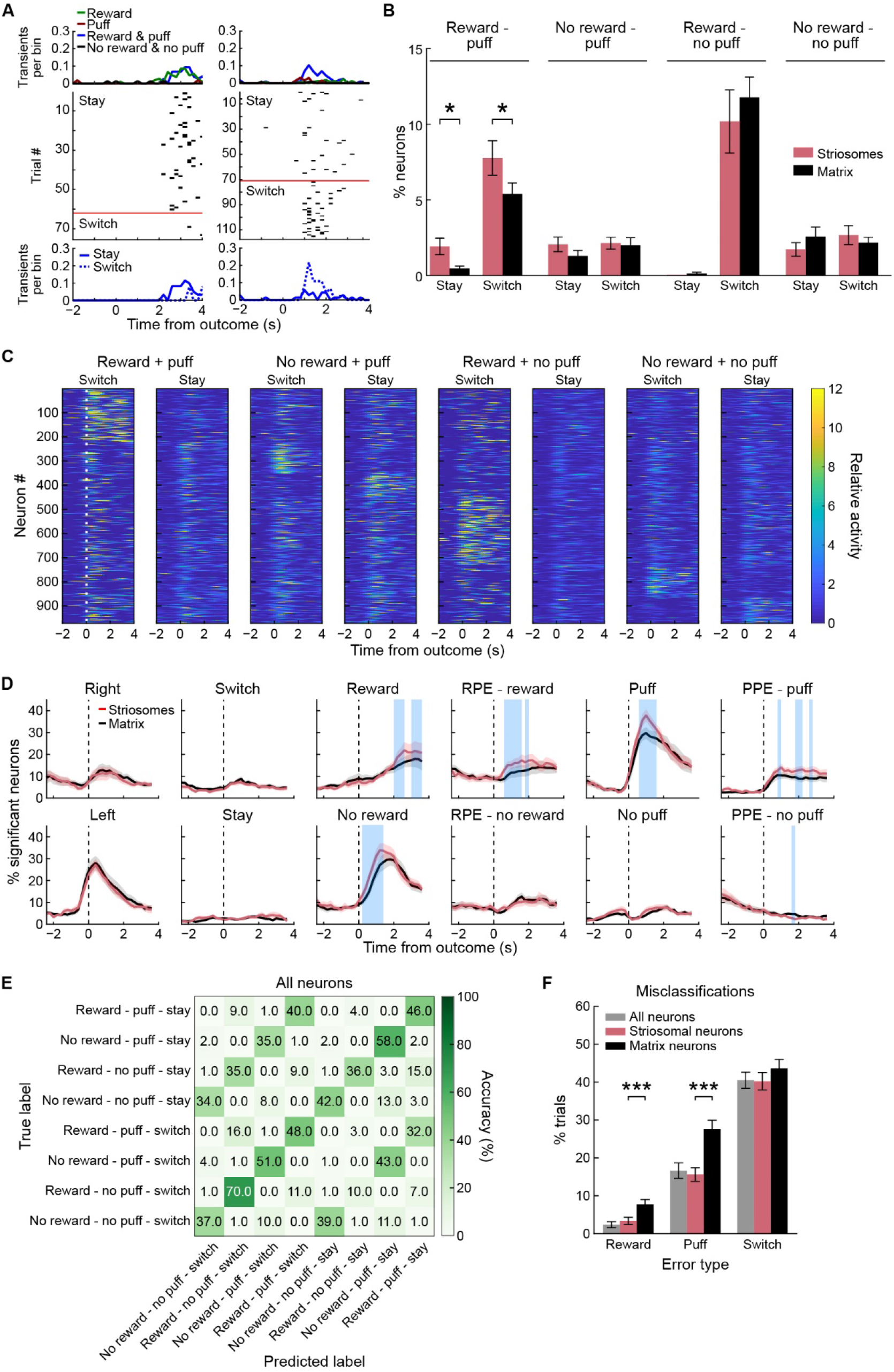
Neuronal Representations of Future Switch/Stay Behavior. (A) Two examples of neurons showing activity in relation to upcoming switch/stay behavior in reward/puff trials. Top: average activity in trials with different reward-puff outcome combinations. Middle: raster plots of all reward-puff trials followed by the same action or the opposite action (above or below the red line, respectively). Bottom: average activity of the neurons in reward-puff trials followed by staying (solid) or switching (dashed). (B) Percentage (mean ± SEM) of neurons that significantly differentiated future staying or switching in trials with different outcome combinations (n = 13 mice). In reward-puff trials, significantly more sSPNs were active in relation to future stay and switch behavior. *p < 0.05. (C) Activity of all neurons showing switch/stay selectivity, averaged per trial type (reward x puff x switch/stay). (D) Time-resolved stepwise regression model showing the average percentage of neurons per mouse that had individual factors included in that timepoint-specific model. Blue shading indicates time points with a significant difference in the percentage of sSPNs and mSPNs (p < 0.05, paired t-test, n = 13 mice). (E) Confusion matrix for decoding of reward-puff-switch/stay trials in a pseudo trial analysis. (F) The sSPN and combined models outperformed the mSPN models when decoding reward or puff outcome but not switch/stay behavior. ***p < 0.001. There were no differences between the sSPN and combined model.

We further assessed the representation of future behavior by regression analyses. First, we identified SPNs encoding reward-puff-switching information (Figure 7C). Second, we performed a time-resolved analysis in which we identified factors predicting activity for every SPN for every time point (Figure 7D). In this analysis, the percentage of neurons with activity representing future switch/stay behavior was low relative to other factors throughout the trial and was not significantly different for sSPNs and mSPNs.

Because we found only weak representations of future switching in single SPNs, we again used ANNs to test whether ensemble SPN activity could predict future switching. Some trial types were extremely rare (e.g., no reward – puff – stay), and so we used only 5 trials per trial type per neurons. We included 51 sessions that had at least 5 trials for all of the 8 reward-puff-switch combinations. Despite the limited samples used for training (32 training trials and 8 test trials), we could decode reward and puff outcomes and predict future switch/stay behavior with a higher accuracy than expected by chance (Figures 7E, S7A and S7B) (misclassifications, reward: 2.4 ± 0.6, puff: 16.6 ± 1.4, switch: 40.5 ± 1.5; Figure 7F). The sSPN model again had lower misclassification rates than the mSPN model for decoding reward (sSPNs: 3.4 ± 0.7%, mSPNs: 7.8 ± 0.9%, p < 0.05) and puff outcome (sSPNs: 15.6 ± 1.2%, mSPNs: 27.6 ± 1.6%, p < 0.05).

Finally, we asked whether SPN activity in the 3 s preceding trial start was predictive of the next action. Models trained with the pseudo trial activity of all SPNs predicted future actions with an accuracy of 85.2%. Models trained with striosomal activity had an accuracy of 86.2%, and matrix models 83.0%. ANN models trained using single sessions had a lower accuracy but one still greater than chance (Figure S7C). Together, these population-decoding analyses demonstrate that the striatal ensemble activity during outcomes and before trial start contained information about the future behavior of the animal.

## DISCUSSION

Our findings in mice demonstrate that single projection neurons in the anterodorsal striatum can represent in their activity associations between an action and both rewarding and punishing outcomes of that action. We observed this action-outcome encoding by the use of a newly developed bandit task in which mice learned without explicit cuing or instruction to maximize beneficial outcome (water) and simultaneously minimize costly outcome (airpuff). In this task, the block sizes deliberately consisted of relatively few trials to force the animals continuously to adapt their behavior in response to evolving prediction errors. Within this CBB task context, we found that visually identified SPNs in striosomes exhibited enhanced encoding of RPE and PPE, relative to nearby matrix SPNs imaged in the same field of view. These findings emphasize multiplexed encoding of action-outcome representations as occurring within the striatum and support a differential function of striosomes in underpinning behavioral adaptation in environments requiring assessment of cost and benefit.

### Multiplexed Encoding of Associations between Actions and Multiple Outcomes by Single Identified Striatal Neurons

We analyzed the Ca^++^ responses of identified SPNs within an RL computational framework. In conventional RL models (Sutton & Barto, 1998), the consequence of an action is usually evaluated by a scalar. Following the concept of RL theories, it is thus tempting to assign the positive value for good and the negative value for bad outcomes by integrating cost and benefit for the evaluative outcome signal. However, when the animals receive multiple modalities for the outcome (i.e., reward and airpuff), neuronal activity can represent the outcome value as a single scalar or can represent the outcomes individually for each modality, as we observed in our sample of SPNs.

The CBB task developed here was an attempt to resolve this ambiguity, as was done previously in humans in a task requiring subjects to maximize monetary rewards and minimize electrical shocks (Seymour et al., 2012). The task allowed mice to learn to associate two different actions with good and bad outcomes and to adapt their behavior rapidly to maximize reward and to minimize punishment. Our findings suggest that in the SPNs imaged, outcomes were not represented as a single scalar value that reflected whether or not an action led to something good. Rather, associations between actions and qualitatively different outcomes were represented in a multiplexed manner in partially overlapping populations of SPNs. We divided SPNs for our analysis into ‘value’ and ‘non-value’ neurons, and found approximately equal numbers of the two. In this task, the outcomes differed both in terms of their valence and their identity, so it was not possible to separate strictly the effect of these two factors on outcome encoding. However, the results do demonstrate unambiguously that outcomes with opposite valence (i.e., reward and punishment) can be represented in parallel, rather in an integrated manner. It remains to be tested whether different outcomes with the same valence are also represented in parallel.

It is possible that the neurons responding in relation to the presence of rewards and puffs, or the absence of both, could signal the overall salience of the outcome. There are two arguments against this explanation. First, we found the outcome activity to be highly action-selective. Second, both delivery and omissions of outcomes could be salient, and we do not observe neurons that were active during salient events, regardless of whether an outcome was delivered or omitted. Because of the extended signaling inherent to Ca^++^ imaging, we also were unable to resolve whether the SPN activity during the decision period reflected the expected outcome or the total expected value. However, the 2-photon imaging preparation that we employed had the great advantage of allowing us to access at a single-cell level the activities of SPNs, and to visually identify their striosomal and matrix identity for over 5000 simultaneously imaged sSPNs and mSPNs.

### Limitations of SPN identification in Targeted Fields of the Anterodorsal Caudoputamen

Pioneering work has shown that direct and indirect pathway neurons (dSPNs and iSPNs) modulate, respectively, approach and avoidance behavior (Kravitz et al., 2012; LeBlanc et al., 2020). Lacking intersectional genetics, we were not able to examine this critical differential representation of outcomes by identified dSPNs and iSPNs. What we could do, however, was to provide a template of information about the attributes of SPNs clustered together in visually identified striosome and matrix compartments.

Visual identification of these SPN subtypes is critical, because genetic models that label striosomes have labeling in the matrix (false positive rate: ∼25%), label striosomes sparsely (false negative rate: ∼70%) and are highly biased to striosomal neurons expressing D1 receptors (Banghart et al., 2015; McGregor et al., 2019; Miyamoto et al., 2018). Our model also suffers from this problem (Bloem et al., 2017; Friedman et al., 2020; Kelly et al., 2018), but the fact that neuropil in the birthdate-labeled striosomes is robustly labeled made it possible to identify visually the striosomes with high reliability (Bloem et al., 2017).

We also emphasize that the results here were all obtained by imaging fields in the anterodorsal striatum, including the region with prominent necklace-like cresent of striosomes (AP range: −0.2-1.2 mm, ML range: 1-2.4 mm); within this region, we did not observe noticeable differences between the way actions, rewards and puff outcomes activated the SPNs, with the caveat that in most mice the range in which we imaged was smaller and we could not directly compare lateral and medial regions, thought to have different encoding properties (Lee et al., 2020; Lerner et al., 2015; Thorn et al., 2010; Yin et al., 2004, 2009).

### Differential Representation of Learning Variables by Visually Identified sSPNs and mSPNs

The spiny neurons within the striosome compartment, sSPNs, are considered to be the main source of striatal input to dopamine-containing neurons of the substantia nigra (SN), whereas mSPNs mainly project to the non-dopaminergic neurons of the SN (Crittenden et al., 2016; Evans et al., 2019; Fujiyama et al., 2011; Gerfen, 1984; McGregor et al., 2019). This parallel innervation has been duly noted: it recalls the parallel network structure of the actor-critic architecture of the RL model (Brown et al., 1999; Doya, 2000; Houk et al., 1995; Joel et al., 2002; O’Doherty et al., 2004; Stephenson-Jones et al., 2013; Suri, 2002; Takahashi et al., 2009). We found that it was in relation to reinforcement-related factors that the striosomal and matrix SPNs differed: sSPNs had more pronounced responses related to both the reward outcomes and to the puff outcomes of given actions, particularly when RPE and PPE were high.

Our findings with this task, indicating that sSPNs are biased to encode RPE and PPE, expand on the previous short reports of the activity of sSPNs and mSPNs in Pavlovian tasks, wherein striosomal activity dominated during reward cues but not in the outcome period (Bloem et al., 2017; Yoshizawa et al., 2018). In Pavlovian tasks, the cues, but not the outcomes, are relevant for updating value expectations in fully trained animals. In bandit tasks, as in the novel CBB task introduced here, it is outcomes, not the trial-start cues, that are critical for updating action values. In this task implementing costs as well as benefits as reinforcers, striosomes and matrix were often co-active, but striosomal activity dominated when new information was provided that resulted in behavioral adaptations, either because of rewarding or aversive consequences. Our findings are thus important conceptually in distinguishing potentially different experimental conditions calling up the activity of striosomes and matrix, as well as in understanding the physiology of potential ‘critic’ circuits.

We did not observe stronger responses related to motor behavior in mSPNs than in sSPNs as often hypothesized. Both during operant, task-relevant behavior and during ITIs, the relationship between neuronal activity and movement was similar in sSPNs and mSPNs. This result at first was surprising, given the strong matrix inputs from the motor cortex and related cortical regions that might access the region from which we recorded (Kincaid & Wilson, 1996; McGregor et al., 2019), but the result is aligned with the view that the striatum is important for learning the value of actions. In our data, we did not observe many neurons that encode motor behavior per se without regard to the outcome.

Our findings for sSPNs and mSPNs suggest that there may be synergistic, cooperative patterns of activation of striosomal and matrix pathways. Such synergism was originally proposed for the D1-expressing direct and D2-expressing indirect pathways (Mink, 1996), and more recent evidence strongly aligns itself with this view of the direct-indirect pathway control system (Tecuapetla et al., 2014, 2016). We suggest that not only the direct-indirect axis of striatal organization, but also the striosome-matrix axis, likely have both synergistic and opposing activity patterns depending on the environmental contingencies requiring adaptive behavioral response.

We draw two central conclusions from this work. First, many outcome-related SPN activities in the dorsal-anterior striatum could not be readily accounted for by a value-coding scheme. Instead, the activities are capable of representing multiple independent action-outcome contingencies (i.e., a multiplex coding). Second, striosomes and matrix exhibit marked differences in their representatons of outcomes, with striosomes biased toward responding in relation to information that is critical for learning processes. These findings provide a new window into how the striatum and its compartmental divisions contribute to adaptive behaviors guided by rewards and punishments in uncertain environments.

## METHODS

### Experimental Model and Subjects

We used 13 Mash1-CreER (het) x Ai14 (het) mice for recording striatal activity, which were offspring of female Mash1-CreER (het) x Ai14 (homo) mice crossed with male C57Bl/6J mice. The female transgenic mice were from a colony that were generated by crossing Mash1(Ascl1)-CreER mice (Kim et al., 2011) (Ascl1tm1.1(Cre/ERT2)Jejo/J, Jackson Laboratory) with Ai14-tdTomato Cre-dependent mice (Madisen et al., 2010) (B6;129S6-Gt(ROSA)26Sor, Jackson Laboratory), which were then crossed with FVB mice to improve breeding results. To generate mice with striosome labeling, we timed the breeding. We paired one male C57Bl/6J mouse with 2 female Mash1-CreER x Ai14 mice. Labeling was induced by injecting pregnant dams with Tamoxifen, dissolved in corn oil, by oral gavage (100 mg/kg) at embryonic day 11.5 (Bloem et al., 2017; Kelly et al., 2018). By this method, Mash1 is expressed predominantly in future striosomal neurons.

We used mice of both sexes (9 females and 4 males) between 6 and 10 weeks old. The experimental animals were housed alone after the first surgery.

### Virus Injections and Surgery

We prepared mice for behavioral training and 2-photon Ca^++^ imaging using previously described procedures (Bloem et al., 2017). Virus was injected in the striatum of adult mice to express GCaMP6s using aseptic stereotaxic surgery. Mice were deeply anesthetized with 3% isoflurane and then head fixed in a stereotaxic frame. Anesthesia was maintained with constant 1-2% isoflurane, and adjusted as needed throughout the surgery. Meloxicam (1 mg/kg) and slow-release buprenorphine (1 mg/kg) were given subcutaneously to provide analgesia. Fur was removed using a depilatory cream (Nair), and the surgical areas were disinfected with three alternating applications of betadine and 70% ethanol. The skin was incised to expose the skull, and the head was leveled to align bregma and lambda in the z-axis. Two burr holes were drilled in the skull, and 500 nl of AAV5-hSyn-GCaMP6s-WPRE-SV40 virus was injected through each hole made at the following coordinates defined relative to bregma: 1) 0.1 mm anterior, 1.9 mm lateral, 2.7 mm ventral; and 2) 0.9 mm anterior, 1.7 mm lateral, and 2.5 mm ventral. Injections were made over 10 min, and the needles was left in place for another 10 min to allow adequate diffusion before retracting. The incision was sutured shut, and mice were kept warm while they recovered from the surgery. Wet food and meloxicam were provided for 3 days to facilitate post-surgical recovery.

Mice were surgically implanted with an imaging cannula at 10-14 days after virus injection. The imaging cannula was assembled by adhering a 2.7 mm glass coverslip to the end of a custom stainless-steel metal tube that was adapted to provide extra stability (1.7 mm long, 2.7 mm diameter eMachineshop; 3D design available at request). Mice were water restricted for a week before the cannula surgery, which significantly improved the clarity of the preparation (Howe & Dombeck, 2016). Mice were deeply anesthetized with isoflurane and head-fixed in the stereotaxic setup. Bregma and lambda were aligned in the z-axis, and craniotomy coordinates were marked at 0.6 mm anterior and 2.1 mm lateral to bregma. The skull was tilted and rolled by 5ᵒ to make it more horizontal at the cannula implant site. A trephine dental drill was used to make a 2.7 mm diameter craniotomy. The cortical tissue was aspirated with gentle suction under constant perfusion with cooled autoclaved 0.01 M phosphate buffered saline until the underlying white matter appeared. A small layer of the white matter was removed and covered with a thin layer of Kwisksil (World Precision Instruments) before inserting the cannula into the cavity. The cannula, as well as a custom head plate, was attached to the skull using metabond (Parkell). Pre- and post-surgical analgesia regimen and care were as described above for the virus surgery.

### Behavioral Apparatus and Task Training

The behavior rig was constructed from optical hardware (Thorlabs) and custom 3D-printed parts (designs available upon request). Mice were head-fixed with their forepaws resting on a Lego wheel, which they could rotate freely, and they reported their decisions by rotating the wheel left or right. The wheel was coupled to a rotary encoder, allowing us to register its rotations with high temporal resolution. The rotary encoder was connected to a microcontroller (Arduino), which ran a custom routine that sampled the position of the rotary encoder every 10 ms. In the event of a movement, the microcontroller sent a timestamp and the amount of movement to a behavioral control computer. The behavioral task was implemented with custom software written in MATLAB (MathWorks) using the Data Acquisition toolbox. Water rewards (3-7 µl, dependent on the behavior of the individual mice) were delivered through a tube by opening a solenoid value for a calibrated period. Air puffs (20 psi) were delivered through a tube positioned on the snout and were similarly controlled with a solenoid valve. Both solenoid valves were located outside the imaging setup. Licking was measure via a conductance-based method (Slotnick, 2009).

We trained mice through successive stages of shaping to implement the final task. Mice were water-restricted for at least one week after recovering from the surgery and received ∼1 ml of water per day. If mice did not earn their water allotment during the task, they were given hydrogel (Clear H_2_O) in their home cage. Mice were first habituated to head-fixation and trained to lick the water tube to receive water rewards. When mice licked reliably, training started. Trial start was signaled with an auditory tone (4 kHz), after which mice had 3 s to move the wheel by 15ᵒ in either direction for it to count as a response. An ITI of 3.5 s was used between trials. To initiate a trial, mice had to hold the wheel still for 2 s. If wheel movement exceeded 5ᵒ in either direction during this time, then an additional 1 s was added to the delay. In the first stage of training, movements to either direction resulted in water reward delivery and no air puff. Reward was delivered when a complete response was registered and signaled with an auditory tone (10 KHz). When mice made responses in at least 150 trials on consecutive days, mice moved on to the next training stage, in which only one action (turning the wheel) was paired with a reward, with contingencies changing after 6-15 rewarded trials. Mice progressed to the next stage of training when they made complete responses in at least 80% of trials, performed ∼200 trials, and showed minimal signs of side bias. In the next stage, air puffs were introduced. In every block, one action was associated with a puff with 100% probability. When mice avoided 6-15 puffs by choosing the action that was not paired with the air puff, the action – air puff contingency switched. Mice progressed to the final task after reaching the same criteria as above.

In the final task, each action was probabilistically (80%) associated with a water reward and an air puff outcome in blocks of trials. For example, in a right puff block, left and right actions were punished with a 0% and 80% probability, respectively. Reward block transition occurred after 6-15 rewards were delivered (chosen randomly for each block). Similarly, puff blocks transitioned after mice avoided 6-15 puffs by selecting the action not associated with puff. Hence, reward and puff block transitions occurred independently so that in some trials the same action was associated with 80% reward and puff probability and in some trials the opposite actions. At the beginning of each session (i.e., the first reward and puff blocks), reward and puff outcomes were independently assigned to one action. At block transitions, the sides associated with reward or puff were switched.

Once mice made responses in at least 150 trials, had no bias and showed clear switch/stay behavior dependent on reward and puff outcomes, they were moved to a behavioral rig under a 2-photon microscope. In most cases, moving from the previous behavioral setup required training the mice for additional 2-7 days until they reached the performance criterion again. The training duration was between 6 weeks and 3 months, with a clear dependence on initial starting age.

### Imaging

Two-photon Ca^++^ imaging procedures were as described previously (Bloem et al., 2017). Briefly, GCaMP6s and tdTomato fluorescence was imaged through a LUMPlan FL, 40x, 0.8NA objective using galvo-galvo scanning with a Prairie Ultima IV 2-photon microscopy system (Bruker). Excitation light at 910 nm was provided by a tunable Ti:Sapphire laser equipped with dispersion compensation (Mai Tai Deep See, Spectra-Physics). Green and red fluorescence emission signals were split with a dichroic mirror (Semrock) and directed to GaAsP photomultiplier tubes (Hamamatsu). Images were acquired at a frame rate of 5 Hz. Laser power at the sample ranged from 11 to 42 mW, depending on the imaging depth and level of GCaMP expression. We selected fields of view (FOVs) that allowed simultaneous imaging of striosomal and matrix neurons. The FOVs had both clearly labeled GCaMPs-expressing cells in striosomes, as defined by dense tdTomato signal in the neuropil, as well as in areas free of tdTomato labeling. Cells were classified as striosomal or matrix depending on whether they were found inside the tdTomato-expressing neuropil zones. In total, we imaged 75 FOVs in 13 mice.

### Image Processing

#### Realignment

We motion-corrected imaging videos by realigning all images to an average reference frame. We first realigned all images in the red stationary channel to the average of all frames in that channel using 2-dimensional cross-correlation (template matching and slice alignment plugin) (Tseng et al., 2011). Next, we realigned the images in the green channel on the basis of the frame-by-frame translations that were calculated for the red channel. We previously found that this procedure does not differentially affect the registration of striosomes and matrix (Bloem et al., 2017).

#### Detection of regions of interest and extraction of DFF

After registration, we detected neurons manually based on the mean, standard deviation and maximum projections. Custom MATLAB scripts were used to calculate local background fluorescence surrounding the somatic regions of interest. The background fluorescence, multiplied by 0.7, was subtracted from the somatic signals, as described previously (Chen et al., 2013). After this, the baseline fluorescence for all neurons (F_0_) was calculated using K-means (KS)-density clustering to estimate the mode of the fluorescence distribution. DFF was calculated as ΔF/F = F_t_ – F_0_ / F_0_.

#### Detection of striosomes

Striosomes were visually identified as regions with dense labeling of tdTomato in the neuropil. We did not record in areas with tdTomato-positive neurons but no clear labeling in the striosomal neuropil. Every cell was classified as striosomal or matrix on the basis of its location in the visually identified compartments. tdTomato-positive neurons outside of the striosomes were included in the striosomal population (32 out of 296 tdTomato-labeled neurons), based on observation that there are exo-patch neurons in the matrix (J. B. Smith et al., 2016) that have similar characteristics as striosomal neurons. Including these neurons or not did not affect our results. In total, 5831 neurons were recorded, of which 2249 neurons were classified as striosomal.

#### Detection of transients

We used a custom algorithm for detecting Ca^++^ transient onsets in the Z-scored DFF. Transients were scored if they met several criteria. First, the size had to be at least 5 times the standard deviation higher than the median DFF. The derivative of the signal also had to exceed the standard deviation, which resulted in detecting onsets. In addition, the mean DFF signal in the 1-s period following the onset, subtracted by the signal in 0.6 s before the onset, had to be bigger than 2 times the standard deviation. After this, in cases where multiple sequential time samples had detected transients, we only kept the first as the event onset. This simple algorithm efficiently detected events (Figure S2A).

### Behavioral Modelling

#### Cost-benefit RL model

We adapted existing RL models (Ito & Doya, 2009; Sutton & Barto, 1998) to include both rewarding and aversive outcomes. We formulated two models. In the first, Q-values for reward and punishment are learned in parallel, and during the decision both are integrated. In the second, there is one set of Q-values, based on the weighed value of an outcome.

##### Parallel RL model

The inferred expectations about reward and puff outcomes linked to actions were modeled using two sets of Q values, one set for reward and one for puffs. In every trial, the decision was modeled using a sigmoid function based on the relative Q-values for reward and the relative Q-values for puff. The sensitivity of the mouse to these differences was parameterized by β_rew_ and β_puff_. A bias term β_0_ was also included.

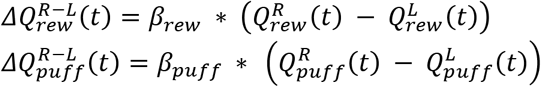

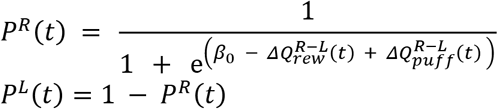

After observing the outcomes, the Q-values for both reward and puff were updated in parallel, using the same rules as in existing models based on reward only. RPE and PPE were calculated by comparing the outcome with the expected value of the chosen action.

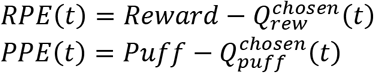

Q-values were updated for both the chosen and the unchosen action. A total of 2 x 3 different learning rates was applied (chosen action followed by outcome: α_rew/puff_; chosen action, no reward/puff delivered α_unrew/nopuff_; non-chosen action: γ_rew/puff_). Bayesian information criterion analysis showed that including all 6 learning rates improved the model.

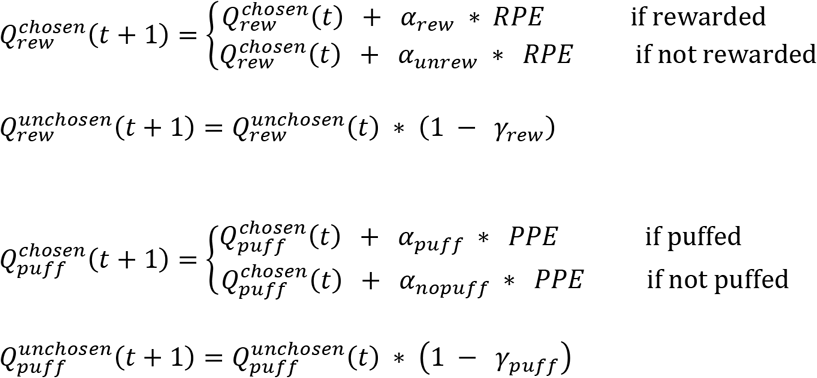

One model was created for every mouse. The model was fit using the fminsearch function in MATLAB. Performance was evaluated using 5-fold cross validation.

##### Integrated RL model

This model is similar to the two set model, but only one set of Q-values is learned. After every trial, the reward and puff outcomes are combined into one outcome value.

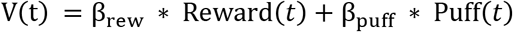

The prediction error is then calculated using this value.

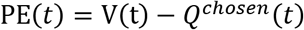

Next, the single set of Q-values is updated.

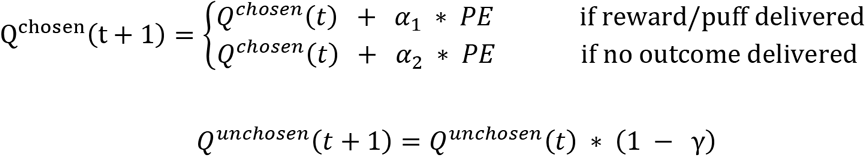

Decision-making is modeled using:

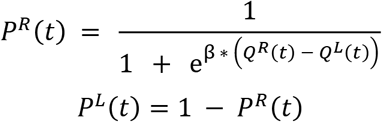

### Auto-regressive behavioral model

We used an autoregressive model to quantify how strongly rewards, puff and actions impact future decision-making (Figure 1G). We fit a model in which we predicted actions in trial *i* with the five previous trials (lagging by *j*). There were 8 sets of predictors. For both left and right (subscript *l/r*), there were *N* (no reward and no puff), *R* (reward, no puff), *P* (puff, no reward) and *B* (both reward and puff). For every trial, one of these was scored as 1 if that particular combination occurred, and the other 7 were scored as 0. A bias term β_0_ was also included. The model was fit for every mouse using the glmfit function in MATLAB. Performance was evaluated using 5-fold cross validation.

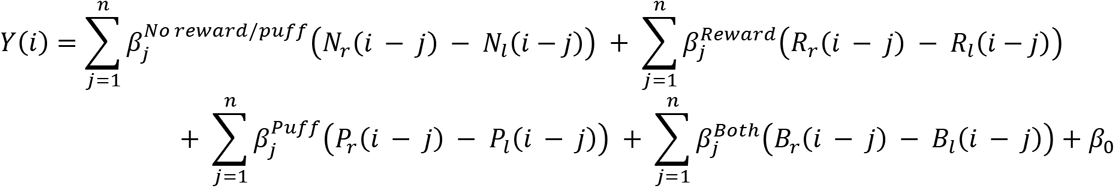

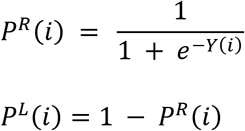

## QUANTIFICATION AND STATISTICAL ANALYSIS

### Analysis of Single-Neuron Activity

#### Analysis of action, reward and puff encoding

To identify neurons that were selective for chosen action, reward or puff outcomes, we used two approaches. First, we used chi-square analysis to test for selectivity by comparing the number of trials with at least one Ca^++^ transient across two conditions (e.g., left vs. right action). To find neurons that were active in relation to multiple features, we performed sequential chi-square tests. First, we tested the individual factors (left vs. right, reward vs. no reward, puff vs. no puff). In neurons that had activity selective for a particular trial type, we then took only these trials, split them again for another factor and ran another chi-square test. Sequential statistical testing is conservative as neurons have to repeatedly pass a statistical criterion. We therefore used a regression as second approach to identify neurons representingactions, outcomes and combinations of these.

We also used a stepwise logistic regression analysis to define neuron types. In these analyses, we selected a number of predictors and then searched for the optimal model for every neuron to predict whether it has a transient in a given trial or not. In every iteration, the effect of adding or removing every possible factor from the model was evaluated. The change that resulted in the largest explanatory power of the model was selected if adding/removing a factor significantly improved the performance of the model. The null hypothesis for the statistical test was that the coefficient of that factor had a coefficient of zero if it would be included in the model. The p-value that was used for cutoff was 0.05. For this analysis, we used the stepwiseglm function in MATLAB.

We performed logistic regression analysis for every neuron to quantify the strength of the relationship between chosen action, reward outcome and puff outcome with the probability of having a transient in a trial. To avoid overfitting, we used L2 regularization.

In all cases, we first calculated the mean per mice and performed statistics on these data. Therefore n = 13 mice for all analyses except stated otherwise.

#### RPE and PPE representations

We analyzed whether neuronal activity in reward/no-reward or puff/no-puff neurons was modulated by RPE and PPE. We used the RL model to infer prediction errors for every trial and then binned these. We then calculated for the different neuronal populations the amplitude of the transients in Z-scored DFF. To test whether neuronal activity was modulated by the prediction errors, we performed correlation analyses. This was done for all trials and also only including the trials with the outcome that the neurons encoded. Statistics were done on the mean activities per mouse (n = 13).

#### Comparing direction selectivity for action-reward and action-puff associations

To test the hypothesis that neurons encode outcome with opposite valence for different actions, we compared the direction selectivity for rewarding and aversive outcomes. First, we used a chi-square test to find neurons that were selective for specific action x reward and action x puff combinations. Then we determined the joint distribution of action x reward and action x puff selective neurons.

#### Selectivity analysis

We compared the selectivity of reward and puff activityin the neurons that were selectively active for both rewards and puff. For this comparison, we calculated for every neuron the proportion of reward/no-reward/puff/no-puff trials in which transients occurred. The selectivity index for reward was calculated by dividing this proportion for trials with a reward by the sum of the proportions of trials with a reward and trials without a reward, resulting in values between 0 and 1, where 1 means that all transients occur in trials with a reward and 0 means that all transients occur in trials without a reward. We did the same for puff selectivity. We transformed these data by subtracting 0.5 and multiplying the outcome by 2 to have a range of −1 to 1, with 0 meaning that transients are as likely to occur in trials with or without the outcome.

#### Wheel movement and licking analysis

We tested how neurons represented wheel movements and licking during the task and in ITIs. Wheel movement and licking data were first binned at 5 Hz to facilitate comparison with the GCaMP6s neuronal signals. For detecting wheel movement bouts, we first smoothed the absolute value of the whole-session movement trace using a 7-point moving average. This facilitated detecting events preceded by ∼1 s of no movement. The resulting trace was baseline adjusted using the MATLAB function ‘ksdensity’ and binarized using a wheel movement threshold of 0.44ᵒ. Event onset and offset times were determined as timepoints at which the binarized movement trace shifted from 0 to 1 and from 1 to 0, respectively. The smoothing introduced a lag of 2 time bins, which was corrected in the onset/offset times. This analysis detects all movement bouts occurring during the session. To restrict analysis to movements occurring during the ITI, we removed all bouts with onset or offset occurring during 3 s from trial start, as well as bouts that had a trial occurring in the middle of them. Individual bouts were labeled as leftward or rightward based on the mean direction of movement. Neuronal data were aligned to time of peak ball acceleration or deceleration within movement bouts. For peak acceleration analysis, significance was determined by comparing the number of movement bouts with neuronal transients occurring in a 1-s window before or after the maximum acceleration using a chi-square test. Significance testing for deceleration was done similarly, except that activity occurring between 0.4 s before and 0.6 s after peak deceleration was compared to a preceding 1-s baseline period.

We performed two types of licking analysis. First, we determined how often licking was coincident with neuronal activity during the ITI period. For each neuron, we generated a peri-stimulus time histogram over a 4-s window by aligning licks to the time of detected neuronal transients. Licks occurring during 3 s after trial start were not counted. Only neurons with at least 10 transients during the session were included in this analysis. We used a shuffle test to generate a null distribution of licking expected by chance. We realigned licking on random permutations of neuronal transient times. This process was repeated 100 times. The total number of licks computed from the observed data was compared to the resulting null distribution. Neurons with observed values outside of the center 95% of the null distribution were considered significant. In a second analysis, we detected licking bouts using a procedure similar to that described for the wheel movements above, with a few differences. The whole-session licking trace was smoothed with a 5-point moving average, and binarization was performed with a lick threshold of 0.2. No baseline adjustment was necessary. For whole-session analysis, neuronal activity was aligned to times of lick bout onsets. For the ITI analysis, only licking bouts occurring outside of trials were used, as described above. Significance testing for both analyses was done by comparing the 1-s pre-licking and 1-s post-licking periods using a chi-square test. For both wheel movement and licking analysis, data from all neurons for individual mice were concatenated together, and the reported values were based on number of mice.

## Decoding Analysis

We used artificial neural networks to decode behavior from the neuronal activity. This resulted in higher accuracy compared to support vector machines or logistic regression and allowed us to decode multiple classes in one analysis without having to create multiple one-versus-rest models. The models consisted of three layers. The input layer used the activity of every neuron. The hidden layer was the same size as the input layer and used ReLU activation functions. The output layer had one node per target and used a softmax function to calculate a probability distribution. We used the Adam optimizer and a learning rate of 0.001, and trained the network in 100 epochs. Dropout (0.3) was used to prevent overfitting. Parameters were chosen on the basis of a grid search.

We created models that were based on pseudo trials and models in which only activity from single sessions was included. We created pseudo trials by selecting from every neuron a fixed number of trials for each target and concatenated across neurons. The number of trials was set separately for the different analysis based on how often trials typically occurred. In the decoding of trial state (action and both outcomes), we used 20 trials resulting in 48 sessions in which all of the 8 trial combinations (chosen action x reward x puff) had more than 20 trials. The other sessions, in which certain combinations did not occur often enough, were excluded. For reward x puff x switch/stay decoding, we used 5 trials per trial type, which was the minimum for 5-fold cross validation and resulted in 51 sessions. For decoding left/right actions based on ITI activity, we included 85 trials, so that all sessions could be included. The pseudo trial analysis was performed 100 times, every time taking different trials from the sessions.

Pseudo trials have the advantage that one can combine neurons from different sessions, which increases the predictive power of the model. However, the number of trials has to be restricted to ensure enough trial samples from all, or most, neurons. In addition, pseudo trials decouple behavioral and neuronal variability. Therefore, we also created models for each session using the same parameters as described above. To compare striosomal and matrix populations, we always took the same number of neurons from every session. The largest population was therefore subsampled. This procedure was repeated 40 times for each analysis.

The models were implemented using TensorFlow (Abadi et al., 2016) in Python 3. Other libraries used were NumPy, SciPy and Pandas.

### Reagent and Resource Sharing

Further information and requests for resources and reagents should be directed to and will be fulfilled by the Lead Contact, Ann Graybiel.

## Data and Code Availability

Code generated in this study will be deposited to GitHub after publication. Data will be available upon request.

## ACKNOWLEDGMENTS

We thank Tomoko Yoshida, Erik D. Nelson and Christian Wuethrich for help with histology, and Dr. Yasuo Kubota for help in preparing the figures and manuscript. This work was supported by NIH/NIMH (R01 MH060379 to A.M.G.; R00 MH112855 to R.H.), Saks Kavanaugh Foundation (to A.M.G.), William N. & Bernice E. Bumpus Foundation (RRDA Pilot: 2013.1 to A.M.G; Postdoctoral Fellowship to B.B.), Simons Foundation (306140 to A.M.G.), Nancy Lurie Marks Family Foundation (to A.M.G.), National Eye Institute (R01 EY028219 and R01 EY007023 to M.S.), National Institute of Neurological Disease and Stroke (U01 NS090473 to M.S.), National Science Foundation (EF1451125 to M.S.), Simons Foundation Autism Research Initiative (to M.S.) and JSPS KAKENHI (20H03555, 20H05469, 20H05063 and 18K19497 to K.A.).

## AUTHOR CONTRIBUTIONS

Conceptualization: B.B, R.H., K.A., and A.M.G.; Software: B.B. and R.H.; Formal Analysis: B.B. and R.H.; Informal Analysis: A.M.G.; Resources: M.S. and A.M.G.; Investigation: B.B., R.H., A.A., G.K., A.W., and C.W.C.; Visualization: B.B. and R.H.; Methodology: B.B., R.H., and A.M.G.; Writing—original draft: B.B., R.H., K.A., and A.M.G.; Writing—review and editing: B.B., R.H., K.A., M.S., and A.M.G.; Funding Acquisition: B.B., R.H., M.S., and A.M.G.; Suprevision: M.S. and A.M.G.

