## Supplementary Figures for "Multiplexed action-outcome representation by striatal striosome-matrix compartments detected with a novel cost-benefit foraging task"

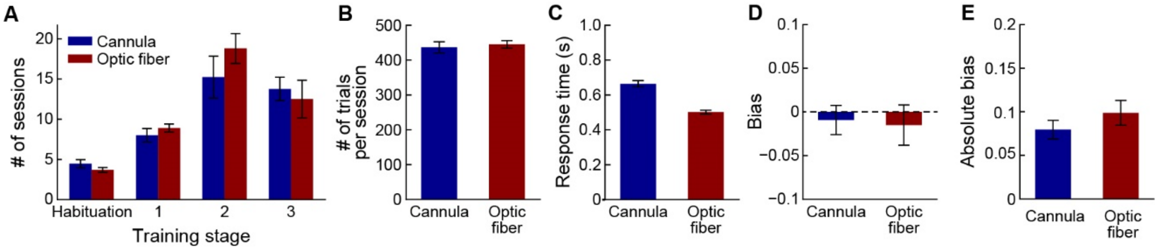


**Figure S1. Effect of Cannula Implantation on Behavioral Performance**

(A) Number of sessions that was required to progress through all training stages, as described in Methods.

(B) Average number of trials performed in the last three sessions before reaching the final performance criterion.

(C) Average response time in the last three sessions before reaching criterion.

(D) Average bias in the last three sessions before reaching criterion.

(E) Average absolute bias in the last three sessions before reaching criterion.

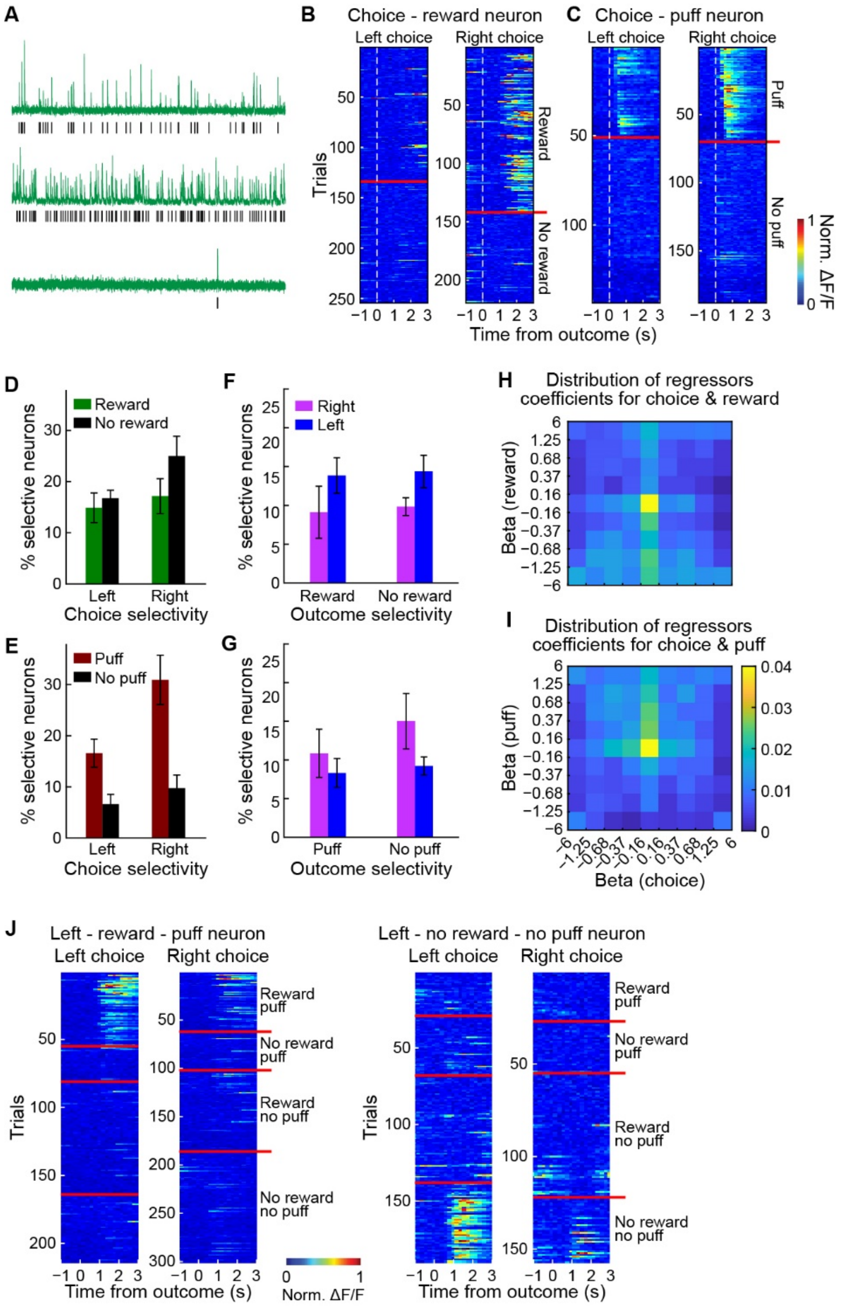


**Figure S2. Action-Outcome Association Representations by SPNs**

(A) DFF fluorescent traces of three sample neurons recorded over 30 min (green) and the time of detected $Ca^{++}$ events (black).

(B and C) Two examples of neurons showing activity selective for action-reward (B) and action-puff (C) combinations. Trials are shown (rows) separately for left/right action, with red lines demarcating reward/no-reward or puff/no-puff outcome trials.

(D) Percentage (mean ± SEM) of action-selective neurons with selectivity for reward or no-reward outcomes (n = 13 mice). There were no significant main effects or interactions.

(E) Percentage of action-selective neurons with selectivity for puff or no-puff trials. There were significant main effects of puff outcome ($p < 0.001$) and choice ($p < 0.05$).

(F) Percentage of reward-outcome-selective neurons with selectivity for left or right actions. No significant main or interaction effects were detected.

(G) Percentage of puff-outcome-selective neurons that was selective for the two actions. No significant effects were detected.

(H and I) Joint distribution of chosen action and reward (H) or puff (I) regressor coefficients. Horizontal and vertical bins were chosen to divide the non-zero coefficients equally among the bins.

(J) Two examples of neurons showing activity representing an association between an action and both reward and puff outcomes.

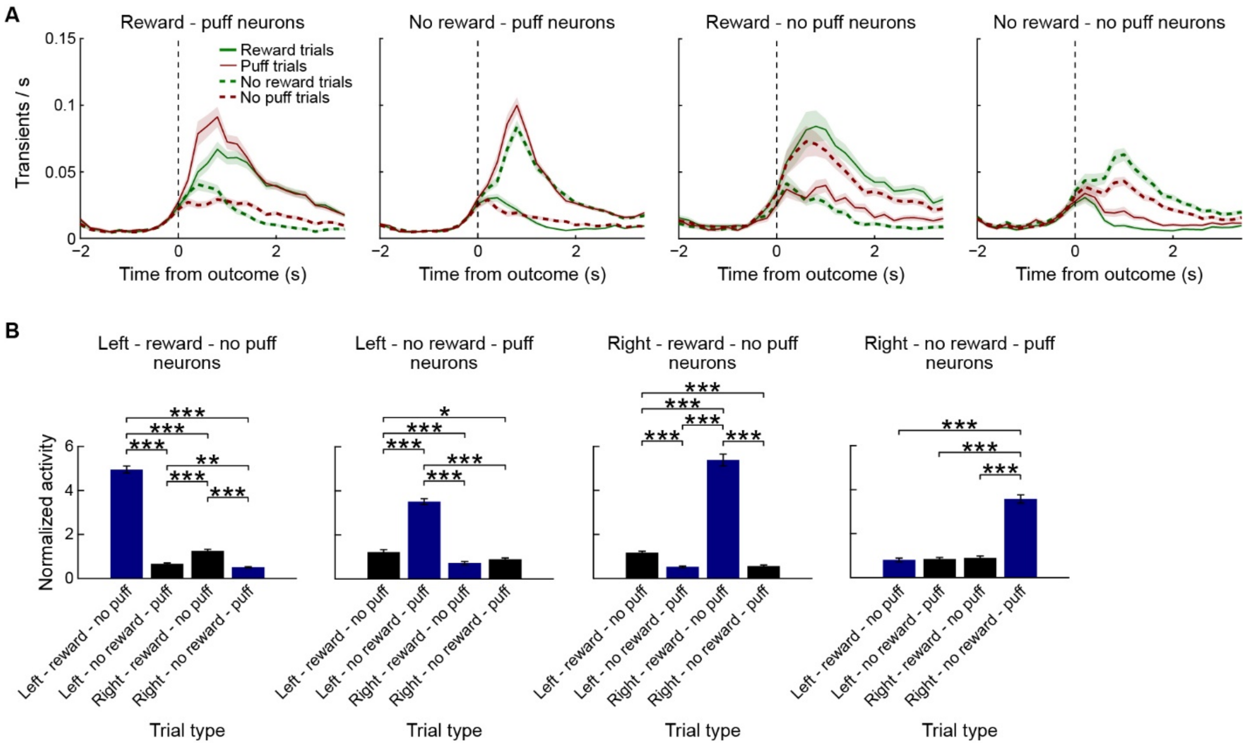


**Figure S3. Activity and Selectivity of Reward, No-Reward/Puff and No-Puff neurons**

(A) Activity (mean ± SEM) of four groups of neurons with activity selective for different outcome combinations in reward/no-reward/puff/no-puff trials (n = 13 mice).

(B) Normalized activity of neurons with value-like responses across different trial types. Neurons that were active in trials in which a good outcome was delivered for one action did not have enhanced activity when the other action was paired with a bad outcome, or vice versa. For all 4 types of neurons, ANOVA indicated significant main effects and interactions ($p < 0.001$). Post-hoc t-test significance levels are indicated with asterisks (*$p < 0.05$; **$p < 0.01$; ***$p < 0.001$).

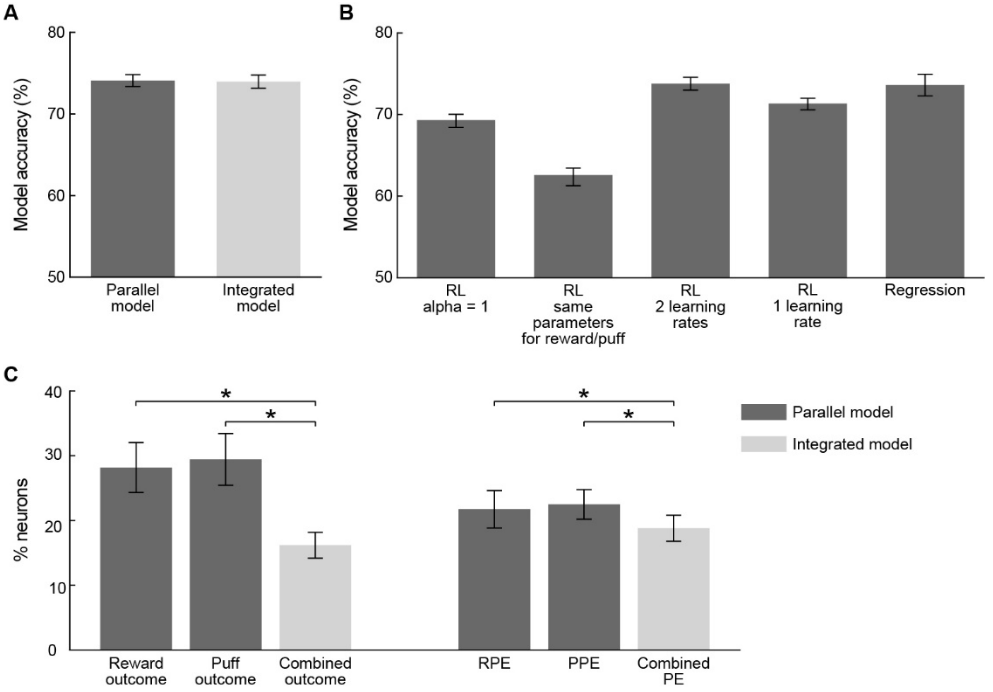


**Figure S4. Comparison of Cost-Benefit Reinforcement Learning Models**

(A) Model accuracy of the parallel and integrated cost-benefit RL models during cross validation.

(B) Cross validation accuracy of alternative simpler models. 'RL alpha = 1': a win-stay/lose-switch model was created by setting all learning rates to 1. 'RL same parameters for reward/puff': one set of learning parameters for both outcomes. 'RL 2 learning rates: the forgetting rate was set to be the same as the unlearning rate (Ito & Doya, 2009). 'RL 1 learning rate: only 1 learning rate was used for each outcome. 'Regression': performance of a 5 trial back auto-regressive model.

(C) A stepwise regression was conducted for each neuron to test which factors best account for the recorded activity. The percentage of neurons that include the factors from the two competing cost-benefitRL models was higher for the parallel model (*$p < 0.05$, average and SEM of 13 mice; paired t-test).

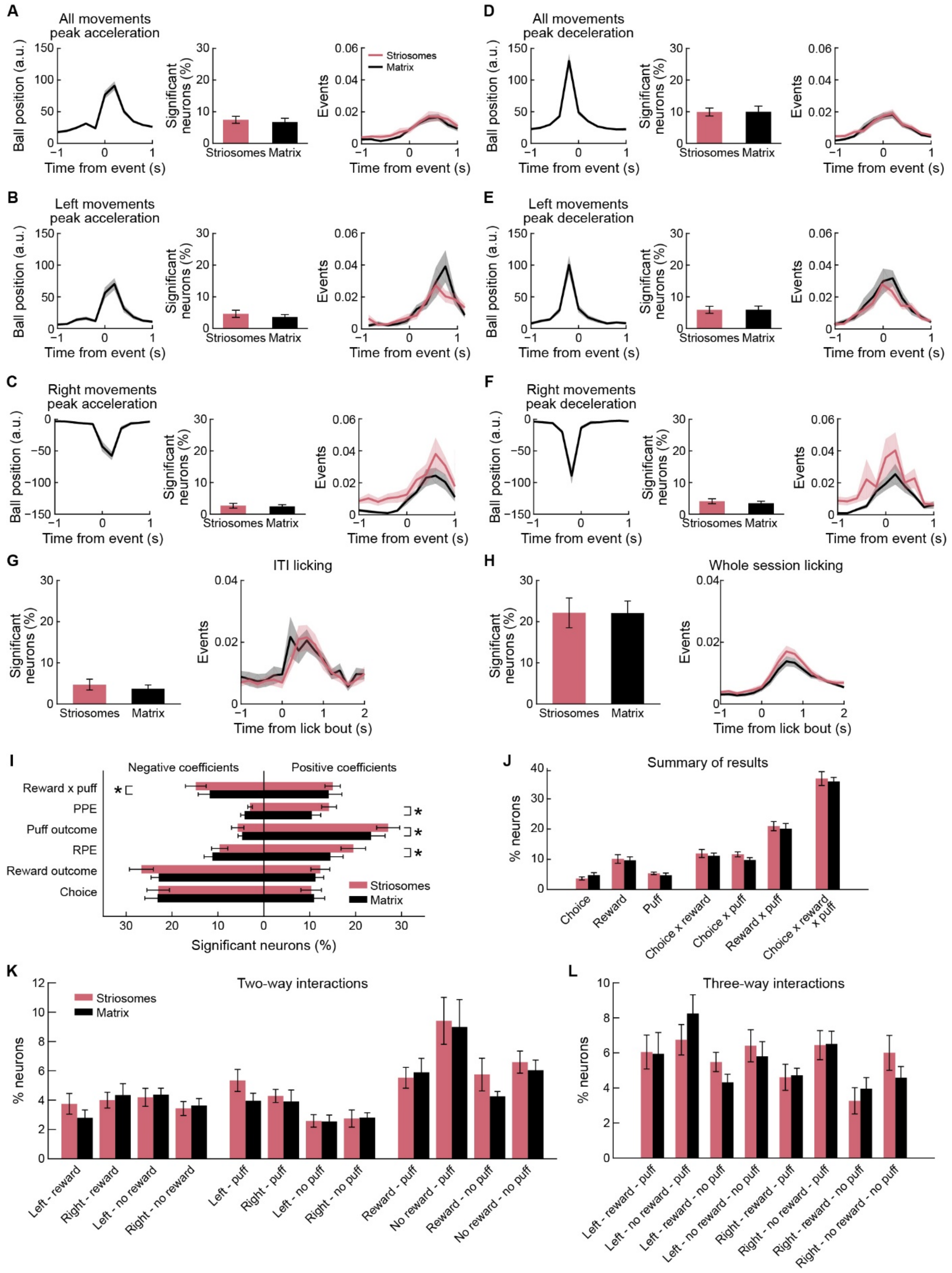
A
All movements peak acceleration
Ball position (a.u.)
Time from event (s)
Significant neurons (%)
Striosomes
Matrix
Events
D
All movements peak deceleration
B
Left movements peak acceleration
E
Left movements peak deceleration
C
Right movements peak acceleration
F
Right movements peak deceleration
G
ITI licking
Time from lick bout (s)
H
Whole session licking
I
Negative coefficients
Positive coefficients
Reward x puff
PPE
Puff outcome
RPE
Reward outcome
Choice
Significant neurons (%)
J
Summary of results
% neurons
Choice
Reward
Puff
Choice x reward
Choice x puff
Reward x puff
Choice x reward x puff
K
Two-way interactions
Left - reward
Right - reward
Left - no reward
Right - no reward
Left - puff
Right - puff
Left - no puff
Right - no puff
Reward - puff
No reward - puff
Reward - no puff
No reward - no puff
L
Three-way interactions
Left - reward - puff
Left - no reward - puff
Left - reward - no puff
Left - no reward - no puff
Right - reward - puff
Right - no reward - puff
Right - reward - no puff
Right - no reward - no puff

**Figure S5. Movement-Related Activity in sSPNs and mSPNs**

(A) sSPN (red) and mSPN (black) activity was aligned to peak acceleration of the absolute value of wheel movement bouts. Panels show average (± SEM) movements across mice (left), proportion of neurons with significant increase in movement-related activity (middle), and mean activity of significantly modulated neurons (right).

(B and C) Similar to A, except movements were divided into left (B) and right (C).

(D-F) Similar to A-C, except activity was aligned to peak deceleration within wheel movement bouts.

(G) Neuronal activity aligned to licking bout onset during ITIs.

(H) Same as G, except for licking bouts occurring during the whole session.

(I) Percentage (mean ± SEM) of sSPNs (red) and mSPNs (black) per mouse that included the chosen action, reward and puff outcomes, their interaction, and RPE and PPE in the optimal model using stepwise regression. Significantly more sSPNs included RPE, puff outcome, PPE and reward x interaction in their optimal model (*$p < 0.05$, $n = 13$ mice).

(J) Summary of stepwise regression showing average percentage of sSPNs and mSPNs per mouse with single action and outcome factors included in their optimal model, as well as different two-way and three-way interactions.

(K and L) Percentage of sSPNs and mSPNs with various two-way (K) and three-way (L) interactions included in their optimal regression model. There are no significant differences in any of the comparisons ($p > 0.05$, $n = 13$ mice).

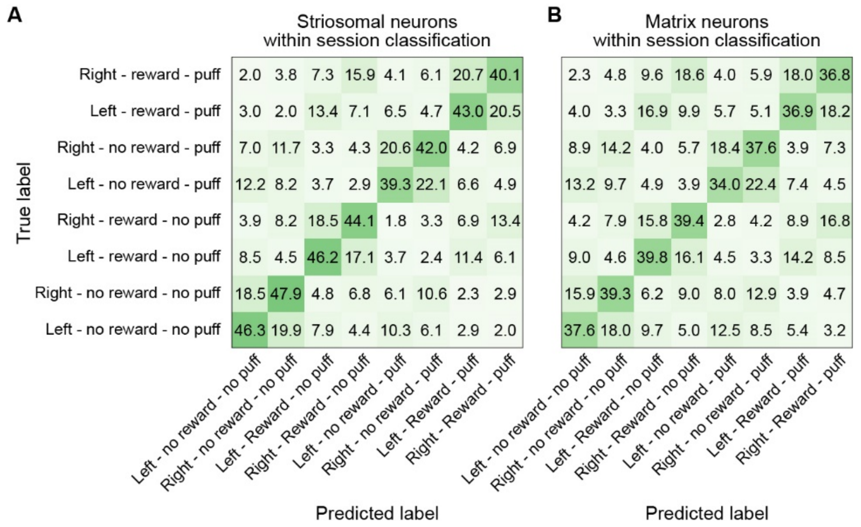


**Figure S6. Decoding Action and Outcome Combinations with Striatal Activity**

(A and B) Confusion matrices for striosomal (A) and matrix (B) decoding of action – reward outcome – puff outcome combinations using single session models.

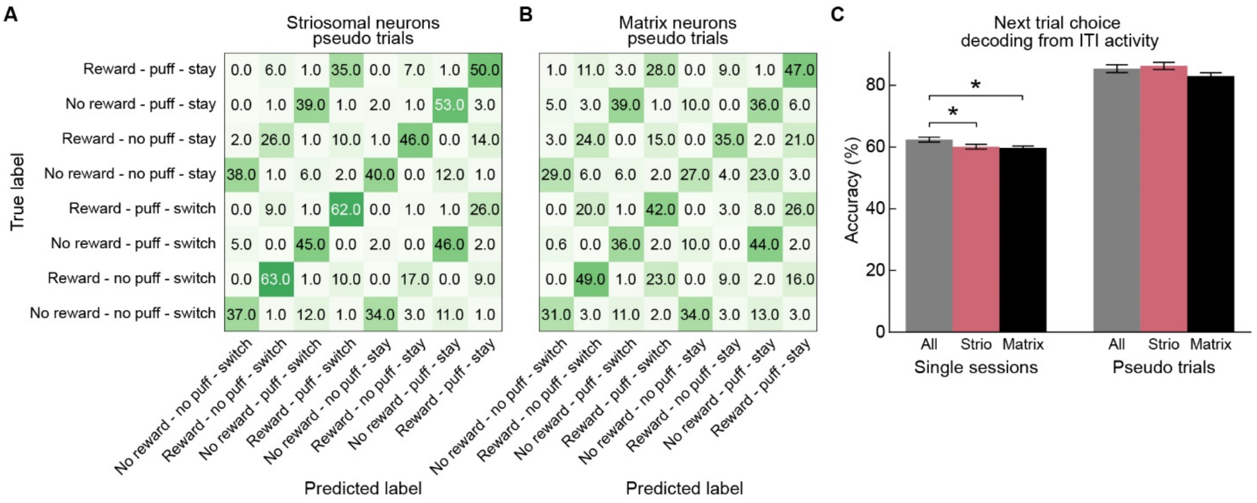


**Figure S7. Decoding Future Behavior with Striatal Activity**

(A and B) Decoding of future switch/stay behavior and reward and puff outcome in striosomes (A) and matrix (B) using pseudo trials.

(C) Accuracy of decoding left/right choices based on ITI activity in the 2 s preceding trial onset (*$p < 0.05$).
